# Directed information flow across the metabolic network of the human brain

**DOI:** 10.64898/2026.03.27.714929

**Authors:** Hamish A. Deery, Chris Moran, Emma Liang, Gary F. Egan, Sharna D. Jamadar

**Author notes:** **Corresponding author address**: Monash Biomedical Imaging, Monash University, 770 Blackburn Rd, Melbourne, 3800, Australia. **e-mail address**.

## Abstract

Brain function is organised in distributed circuits in which regional engagement unfolds over time, reflecting coordinated and temporally ordered patterns of neural computation and information flow. Neural activity also depends on a reliable and scalable supply of glucose. Yet, the temporal order and direction of metabolic signalling in brain circuits remain unknown. Here, we combine functional Positron Emission Tomography (fPET) with [18^F^]-flurodeoxyglucose and Granger causality analysis to characterise directed metabolic connectivity in cognitive control, memory and affective regulatory circuits in 86 healthy adults. We observed widespread directed metabolic influences within the circuits, with the strength of connections a significant predictor of cognition and affect. The behavioural value of the connections was also governed by the efficiency with which baseline glucose metabolism was converted into adaptive functional connections. We conclude that the brain is organised into metabolic circuits that coordinate temporally ordered connectivity to support information transfer. Directed connections vary in their efficiency of glucose use and functional benefit, suggesting that metabolic signalling does not follow a simple “more is better” rule but reflects context-dependent optimisation across cognitive systems.

**Author Summary:** Activity in the regions of brain circuits unfolds over time, reflecting coordinated and temporally ordered patterns of neural computation. However, it is currently unclear how the intrinsic architecture of the human brain’s metabolic network supports temporally ordered information flow. Using functional Positron Emission Tomography (fPET) and Granger causality analysis, we identify widespread directed metabolic influences within cognitive control, memory, and affective regulatory circuits. We report that metabolic signalling reflects structured, hierarchical dynamics that predict cognitive performance and psychosocial function. Our results indicate that the brain optimises energetic resources through context-dependent signalling: proactive control requires high metabolic investment, while memory circuits benefit from economical use of glucose. These findings suggest that the brain’s metabolic architecture is organised to support information flow in functional circuits.

## Introduction

All our thoughts, feelings and behaviours emerge from coordinated and ordered patterns of neuronal activity across distributed functional brain networks (Bullmore & Sporns, 2009; Liegeois et al., 2017; Singer, 2009; Wig, 2017). Coherent fluctuations in neuronal activity within and between networks are thought to be the mechanism behind information transfer in the brain, and therefore, cognition and psychosocial function (Edelman, 1993). The functional brain circuitry that supports cognition depends upon a *reliable* and *scalable* (i.e., able to be up– and down-regulated on demand) supply of glucose, the brain’s primary energy substrate (Jamadar et al., 2025; Mergenthaler et al., 2013). Coherent *glucodynamic* activity between distributed brain regions forms the metabolic network of the brain (Deery, Liang, Siddiqui, et al., 2024; Jamadar, Ward, et al., 2021), and interactions within and between metabolic sub-networks influences cognitive performance (Voigt et al., 2023). Normative changes in cerebral glucose metabolism and the metabolic network across the lifespan are associated with age-related cognitive decline (Deery et al., 2023; Deery, Liang, Siddiqui, et al., 2024). Here, we explore how information flows directionally from one region of the metabolic network to another, and how this influences cognition and psychosocial function and is altered in ageing.

Activity in the regions of brain circuits unfolds over time, reflecting coordinated and temporally ordered patterns of neural computation. The synaptic activity, integrative processing, and sustained firing of neural populations that occurs during neural computation is energetically demanding, and is associated with an increase in cerebral glucose uptake and metabolism at the post-synaptic terminal (Howarth et al., 2012; Kety, 1957; Sokoloff, 1960). The dynamics with which synaptic glucose metabolism varies over time is an important determinant of cognition (Deery et al., 2026; Jamadar et al., 2025). A larger dynamic range and complexity of glucodynamic activity is associated with better cognition, enabling the brain to transfer information efficiently and enter a rich range of network states (Deery et al., 2026; Deery, 2025). However, it is currently unknown how information flows across the metabolic network, and whether metabolic activity in one region precedes and predicts subsequent activity in another. Within this framework, temporal dependencies in regional glucose metabolism can be considered a marker of directed information transfer that captures the order in which regions contribute to ongoing processing. For example, when applied to cognitive control circuits, this perspective predicts that metabolically demanding control regions, such as the dorsolateral prefrontal cortex, should exhibit earlier or predictive metabolic activity relative to downstream regions involved in action selection or execution, consistent with hierarchical models in which control signals bias and sustain processing across the circuit (Braver, 2012; Ruge et al., 2013). Directed analyses of time-variant glucose signals therefore offer a principled means of testing hypotheses about the flow of information processing in brain circuits.

Traditionally, neuroimaging studies have used haemodynamic (fMRI) and electrophysiological (EEG/MEG) signals to characterise neuronal activity and the network architecture of the brain. Mechanistic approaches, such as Granger causality analysis (Ding et al., 2000; Goebel et al., 2003; Granger, 1969) and dynamic causal modelling (DCM (David et al., 2006; Friston et al., 2003)), have been used to infer directed influences between brain regions, offering insight into the causal architecture of circuits underlying brain-behaviour relationships (Friston et al., 2013; Reid et al., 2019). Together, these approaches have revealed directed information flow across electrophysiological and haemodynamic circuits that support cognitive function, such as cognitive control (Braver, 2012; Kok et al., 2006) and memory (Baddeley, 2003; Constantinidis & Procyk, 2004; Moore et al., 2013), as well as circuits that regulate psychosocial function and underlie anxiety (Andreescu et al., 2015; Coutinho et al., 2016) and depression (Liu et al., 2023; Zhong et al., 2011) symptomology. It is unknown whether the information flow in these circuits is sustained by similar directional metabolic signal.

Examination of directed connectivity can be categorised as *model-based and model-free* (Seth et al., 2015). DCM is the most commonly used model-based approach, applying a theoretically and mathematically defined biophysical model to understand directed *effective* connectivity (Friston et al., 2014). By comparison, the model-free approach of Granger causality does not invoke a biophysical model; rather, it applies a linear vector autoregressive (VAR) model to infer that activity in brain region X ‘Granger-causes’ activity in brain region Y (Seth et al., 2015). According to this model, if the preceding activity in X predicts the future activity of Y, above and beyond the information in the timeseries of Y, then it is inferred that information has ‘flowed’ from region X to Y (Seth et al., 2015). The fact that Granger causality does not rely on a defined mathematical model means that it is well-suited to understand directed interactions in data for which such models do not exist, such as for glucodynamic data.

Here, we use Granger causality analysis as a modelling framework for directed metabolic connectivity. Cerebral glucose uptake is measured using [18^F^]-fluorodeoxyglucose positron emission tomography (FDG-PET) (Clarke, 1999; Sokoloff, 1960; Ter-Pogossian, 1992), with time-variant glucodynamics measured with the ‘functional’ PET (fPET) approach (Deery et al., 2026; Deery, Liang, Siddiqui, et al., 2024; Hahn et al., 2020; Hahn et al., 2016; Hahn et al., 2024; Jamadar, Ward, et al., 2021; Rischka et al., 2018). We extract glucodynamic timeseries from regions involved in cognitive control, working and episodic memory, and affective regulation, and apply Granger causality analysis to investigate their directed interactions and associations with cognition and psychosocial function.

Guided by theoretical models of cognition and prior neuroimaging work, we test four hypotheses. First, we hypothesise that directed metabolic connections will reflect hierarchical organisation in cognitive control circuits, with strong top-down connections from prefrontal cortical regions to downstream cortical and subcortical targets. Second, we hypothesise that working memory will depend on efficient frontoparietal information transfer, particularly in attention-control pathways. Third, we hypothesise that episodic memory will depend on hippocampal-driven information transfer to cortical regions. Finally, we hypothesise that individual differences in anxiety and depression will relate to directed metabolic signalling in frontolimbic and salience circuits.

## Results

We used Granger causality analysis of fPET data from 86 healthy adults aged 20-86 years (see Supplement 2 for demographics) to examine directed metabolic connections between brain regions in theoretically motivated circuits for cognitive control, working and episodic memory and affective regulation. The ROI coordinates and associated Brodmann areas, as well as the literature on the connections tested in the circuits, are described in detail in Supplement 3. They are also described briefly below.

All analyses were conducted separately within each hemisphere (with midline structures modelled centrally), as no *a priori* hypotheses were specified regarding laterality or interhemispheric metabolic connectivity. For each predefined source-target connection, the directed metabolic influence was quantified using a vector autoregressive (VAR) model with a lag of two fPET frames (see Methods for discussion of model parameter choices, VAR equations, and multi-lag sensitivity analysis). We first characterised directed metabolic connectivity in each circuit and test the connections for statistical significance compared to zero and a biologically plausible null model using circular temporal shifting and false discover rate (FDR) correction. We then used correlation analyses to test associations between the directed connections and cognitive performance or affective symptom measures corresponding to each circuit’s functional domain. We also tested for age group differences in these associations.

We also undertook exploratory analyses to evaluate the baseline energy use and the metabolic efficiency of circuit-level directed metabolic signalling. We quantified the cerebral metabolic rate of glucose (CMR_GLC_) and a “glucose cost index” (GCI; the product of the Granger and the CMR_GLC_ values of the constituent ROIs; see Methods for equation) for directed connections. We defined CMR_GLC_ as the baseline metabolic budget – the total energy consumed by the constituent regions – while the GCI represents the metabolic investment required to sustain communication between them. By distinguishing between these measures, we assess whether a connection’s significant association with cognition is driven by a high baseline budget (CMR_GLC_) or by the efficiency with which that energy is converted into a behavioural outcome (GCI). For instance, a behaviourally relevant connection characterised by a high baseline CMR_GLC_ but a low GCI indicates an efficient pathway, where substantial metabolic resources are managed with minimal processing cost, suggests efficient utilisation of a relatively high glucose budget.

### Cognitive control circuit

For the cognitive control circuit, we defined divisions based on the dual mechanism of control model (Braver, 2012) and used ROIs based on neuroimaging studies of task switching (measuring proactive control) and response inhibition (reactive control) (Braver et al., 2003; Emch et al., 2019; Hughes et al., 2014; Jamadar et al., 2010; Kok et al., 2006; Marklund & Persson, 2012; Ruge et al., 2013; Yin et al., 2015). The proactive control circuit comprised the dorsolateral prefrontal cortex, anterior cingulate cortex, and caudate, with directed pathways reflecting sustained goal maintenance and cortico-striatal information transfer (Figure 1a). The reactive control circuit comprised the anterior insula, ventrolateral prefrontal cortex, inferior frontal junction, and superior parietal lobule, with directed influences measuring information transfer in control and attention pathways (Figure 1b). Granger causality analysis revealed directed metabolic influence in the proactive and reactive control circuits (Figure 1a-b; full statistics in Supplementary Table S6). All connections were significantly different to zero (p-FDR < 0.001). When compared to the permutation null model, several of the strongest pathways showed significantly greater metabolic influence than expected by chance. In the reactive control circuit, the right inferior frontal junction to the superior parietal lobule connection was significant (mean GC = 0.019, p-FDR = 0.036). In the left hemisphere of the proactive control circuit, connections from the dorsolateral prefrontal cortex to the caudate (mean GC = 0.019, p = 0.013) and the anterior cingulate to the left caudate (mean GC = 0.018, p = 0.022) reached nominal significance but did not survive FDR-correction (both p-FDR = 0.099). Collectively, these results indicate widespread metabolic Granger causality in the cognitive control circuits, with a subset of connections representing especially salient and above-chance metabolic signalling pathways. The finding of left hemisphere predominance for proactive control and right predominance for reactive control is consistent with classic models of response selection and inhibition in frontoparietal circuits (Aron & Poldrack, 2006; Jamadar et al., 2010; Ridderinkhof et al., 2004).

**Figure 1.**
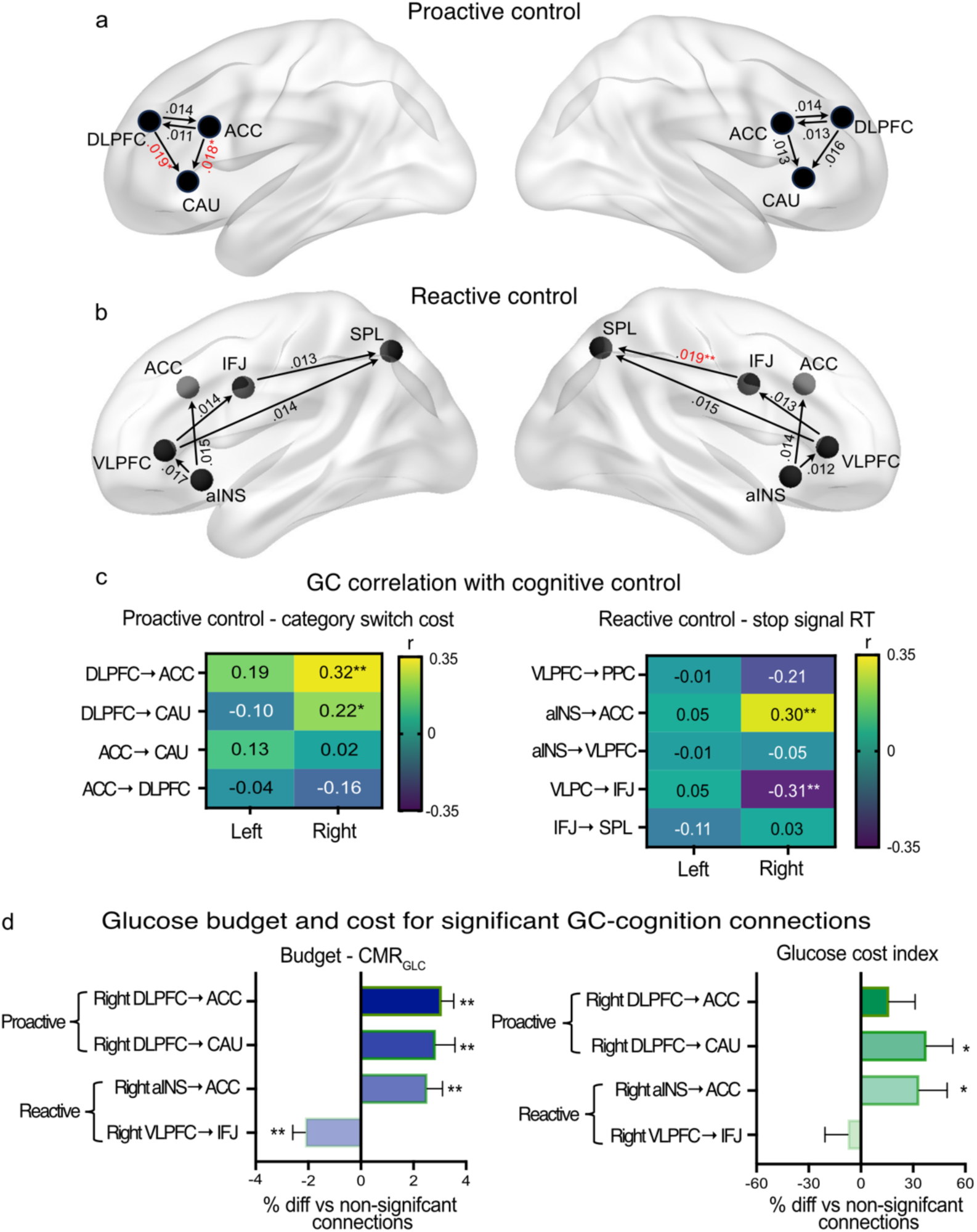
Directed metabolic connectivity in the cognitive control circuit. Granger causality results in the cognitive control circuit for proactive (a) and reactive (b) control divisions. (c) Correlations between Granger causality values and category switch and stop signal task performance. (d) Mean percentage difference in glucose budget (CMR_GLC_) and glucose cost index of connections with cognitive performance associations versus non-significant connections (error bars are standard error of the mean). *p < 0.05; **p-FDR < 0.05 (in panel a, p-values are compared to a permutation null model). ACC = anterior cingulate cortex, corresponds to Brodmann area (BA) 32; aINS = anterior insula (BA13); CAU = caudate; DLPFC = dorsolateral prefrontal cortex (BA9/46); IFJ = inferior frontal junction (BA6/44); SPL = superior parietal lobule (BA7); VLPFC = ventrolateral prefrontal cortex (BA44/45/47). Note that for visualisation purposes only, the location of some ROI nodes in panels a-b have been modified slightly from their original MNI coordinates to enable the pathways to be clearly displayed.

To test our hypothesis that directed metabolic connections will reflect hierarchical organisation in the cognitive control circuits, we correlated GC values in the proactive and reactive divisions with switch cost and stop signal reaction time, respectively (Figure 1c). The cognitive measures were multiplied by –1 so that higher switch cost indicates better cognitive flexibility, and higher stop signal reaction time indicates faster response inhibition. For proactive control, stronger connectivity from the right dorsolateral prefrontal cortex to the anterior cingulate (r = 0.32, p = 0.002, p-FDR = 0.010) was associated with better cognitive flexibility (full statistics in Supplementary Table S7). Similarly, stronger connectivity from the right anterior insula to the cingulate predicted faster response inhibition (r = 0.30, p = 0.005, p-FDR = 0.012). In contrast, stronger right ventrolateral prefrontal cortex to inferior frontal junction connectivity was associated with slower response inhibition (r = –0.31, p = 0.005, p-FDR = 0.012). These results indicate that in some instances, information flow in metabolic circuits may actually interfere with cognitive performance. To more fully explore these results, Supplement 4.2 presents *post hoc* analyses, the results of which indicate that activity within and between the circuits are additive; that is, connections in one part of a circuit do not ameliorate interference effects in another.

We now turn to the exploratory glucose budget (CMR_GLC_) and GCI analyses to test whether behaviourally relevant connections have different levels of metabolic supply and efficiency compared with connections without behavioural relevance. We computed a percentage difference for each subject for each ROI for CMR_GLC_ and GCI. First, for the proactive and reactive control circuits separately, we calculated the per-subject percentage difference between each significant and all non-significant behavioural connections to expresses the metabolic cost of the significant connections. We then tested whether the mean percentage difference between significant and non-significant connections across subjects differed from zero using a one-sample *t*-test, thresholded at α=0.05 for these CMR_GLC_ and GCI exploratory analyses (Figure 1c).

Connections identified as behaviourally relevant in the right proactive control circuit (DLPFC → ACC and DLPFC → CAU) exhibited higher baseline CMR_GLC_ of 2.9% and 2.4% compared with non-behaviourally relevant connections (p < 0.001; Figure 1d). Similarly, in the reactive control circuit, the right anterior insula to cingulate connection showed 2.1% higher CMR_GLC_ (p < 0.001). In contrast, the only connection showing a negative relationship with behaviour (VLPFC → IFJ), demonstrated a 2.0% *lower* CMR_GLC_ (p < 0.001). In terms of the GCI, significant behaviourally relevant connections also exhibited greater glucose cost than non-significant connections, with right dorsolateral prefrontal cortex to caudate and anterior insula to cingulate showing increases of 38% and 33% (p = 0.018 and p = 0.046).

Together, these results indicate that the brain differentially adjusts glucose metabolism and information transfer across proactive and reactive control pathways, balancing efficiency of glucose use and signal strength to optimise cognitive performance depending on the type of control required. Behaviourally relevant connections in the proactive division have a higher baseline glucose budget, with dorsolateral prefrontal to cingulate pathways operating efficiently and dorsolateral to caudate connections showing both high cost and strong directed influence. By contrast, connections in the reactive control circuit display distinct patterns: right anterior insula to cingulate connections combine high metabolic cost with directional influence, supporting high cost, high yield reactive processing, whereas ventrolateral to inferior frontal junction connections have a lower baseline glucose budget and negative behavioural associations, suggesting low metabolic investment for poor functional return.

### Working memory circuit

The working memory circuit was divided into attention-control and storage-maintenance components (Hartley & Speer, 2000; Liu et al., 2025) consistent with the literature (also see Supplement 3). The attention-control division included the dorsolateral prefrontal cortex, intraparietal sulcus, and medial frontal cortex, supporting executive manipulation, attentional shifting and top-down monitoring. The storage-maintenance division comprised the ventrolateral prefrontal cortex and inferior parietal lobule, supporting rehearsal and phonological storage. Directed pathways modelled the information transfer between executive control and stored representations (Figure 2a-b). Analysis of the working memory circuit revealed widespread directed metabolic flow, with all individual Granger causality values significantly greater than zero (p-FDR < 0.05, one-sample *t*-tests). The largest absolute deviations were observed along the left medial frontal cortex to inferior parietal lobule (mean GC = 0.018) and the right dorsolateral to ventrolateral prefrontal cortex (mean GC = 0.018) pathways. However, when evaluated against the more conservative permutation null model, these pathways did not survive false discovery rate correction, indicating that while the circuit exhibits global directed signalling, these specific regional paths should be interpreted as trends.

**Figure 2.**
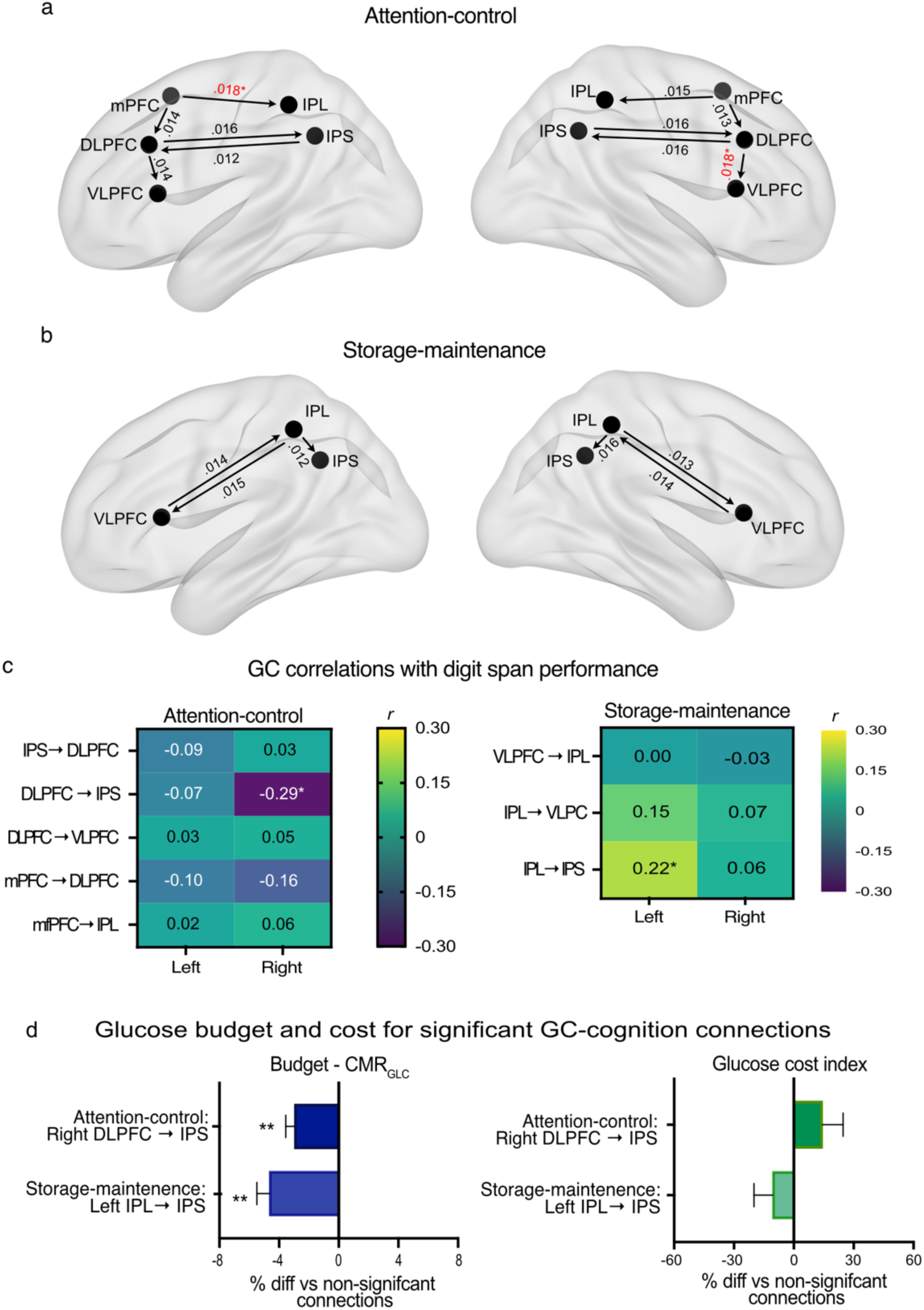
Directed metabolic connectivity in the working memory circuit. Granger causality results in the working memory circuit for the attention-control (a) and storage-maintenance (b) divisions. (c) Correlations between Granger causality values and working memory performance. (d) mean percentage difference in glucose budget (CMR_GLC_) and glucose cost index for connections with cognitive performance associations versus non-significant connections (error bars are standard error of the mean). *p < 0.05; **p-FDR < 0.05 (in panel a, p-values are compared to a permutation null model). VLPFC = ventrolateral prefrontal cortex (BA 45/47), IPL = inferior parietal lobule, IPS = intraparietal sulcus, DLPFC = dorsolateral prefrontal cortex (BA 9/46); mPFC = medial frontal cortex (BA 6/32). Note that for visualisation purposes only, the location of some ROI nodes in panel a-b have been modified slightly from their original MNI coordinates to enable the pathways to be clearly displayed.

To test our hypothesis that working memory will depend on frontoparietal information transfer, we correlated the GC values with the total longest forward and backward digit span, with higher scores indicating better performance. These analyses revealed moderate strength, lateralised patterns of association (Figure 2c). Stronger connectivity from the right dorsolateral prefrontal cortex to inferior parietal sulcus in the attention-control circuitry was associated with lower working memory capacity (r = –0.29, p = 0.007, p-FDR = 0.055). In contrast, stronger connectivity from the left inferior parietal lobule to intraparietal sulcus in the storage-maintenance circuit showed a trend towards higher working memory capacity (r = 0.22, p = 0.047, p-FDR = 0.376). Therefore, metabolic connections in this circuit are at times beneficial for memory performance, and at other times, appear to have a moderately detrimental impact on performance. Supplement 4.2 reports a *post hoc* analysis to further explore this result, which showed the significant connections to be additive rather than compensatory in their associations with memory performance.

In terms of the glucose budget and cost analyses, connections that showed memory performance relevance had lower CMR_GLC_ compared with non-significant connections (Figure 2d). Specifically, the connections between the right dorsal prefrontal cortex and intraparietal sulcus in the attention-control circuit had a 3.2% lower CMR_GLC_ (p < 0.001), suggesting that effective working memory-related signalling may occur with a comparatively low glucose budget. Similarly, the connection between the left inferior parietal lobule and intraparietal sulcus in the storage-maintenance circuit had a 5.2% lower CMR_GLC_ than non-significant connections (p < 0.001). When considering the GCI, the left inferior parietal lobule to intraparietal sulcus and the dorsolateral prefrontal cortex to intraparietal sulcus pathways showed no difference in GCI to non-significant connections (p = 0.272 and p = 0.173), suggesting no difference in cost-weighted directional influence despite lower baseline glucose budgets.

Together, these results suggest that directed metabolic signalling in the working memory circuit is organised by a functional division between storage-maintenance and attention-control, with distinct implications for information transfer. Connections within the storage-maintenance division support working memory with a low glucose budget, indicating that sustained representational processes rely on efficient, low-cost directional communication. In contrast, effective working memory-related signalling may occur with a comparatively low glucose budget. Overall, the findings imply that effective working memory depends on the selective routing of information between posterior storage and anterior control regions, where optimal performance reflects a balance between metabolically economical maintenance signalling and top-down control of information transfer.

### Verbal episodic memory circuit

The verbal episodic memory circuit was divided into encoding-binding and reconstruction-retrieval divisions (Thommesen et al., 2024) based on the literature (also see Supplement 3). The encoding-binding circuit comprised the ventrolateral prefrontal cortex, perirhinal cortex and hippocampus, with directed influences measuring memory encoding and relational binding in medial temporal lobe structures. The reconstruction-retrieval system included the posterior middle temporal gyrus, angular gyrus, hippocampus, perirhinal cortex and ventrolateral prefrontal cortex. Directed pathways modelled item– and relational-level reinstatement, integration within parietal structures and feedback to prefrontal control systems during retrieval monitoring (Figure 3a-b). Directed metabolic influence was observed across the verbal episodic memory circuit, with all connections significantly different from zero (p-FDR < 0.05, one sample *t*-tests). The strongest pathway was from right ventrolateral prefrontal cortex to perirhinal cortex (mean GC = 0.017). When tested against the permutation null model, this pathway did not survive FDR correction (p = 0.022; p-FDR = 0.198). Hence, like the working memory circuit, the episodic memory circuit exhibits global directed signalling across connections, although specific paths should be interpreted as descriptive trends.

**Figure 3.**
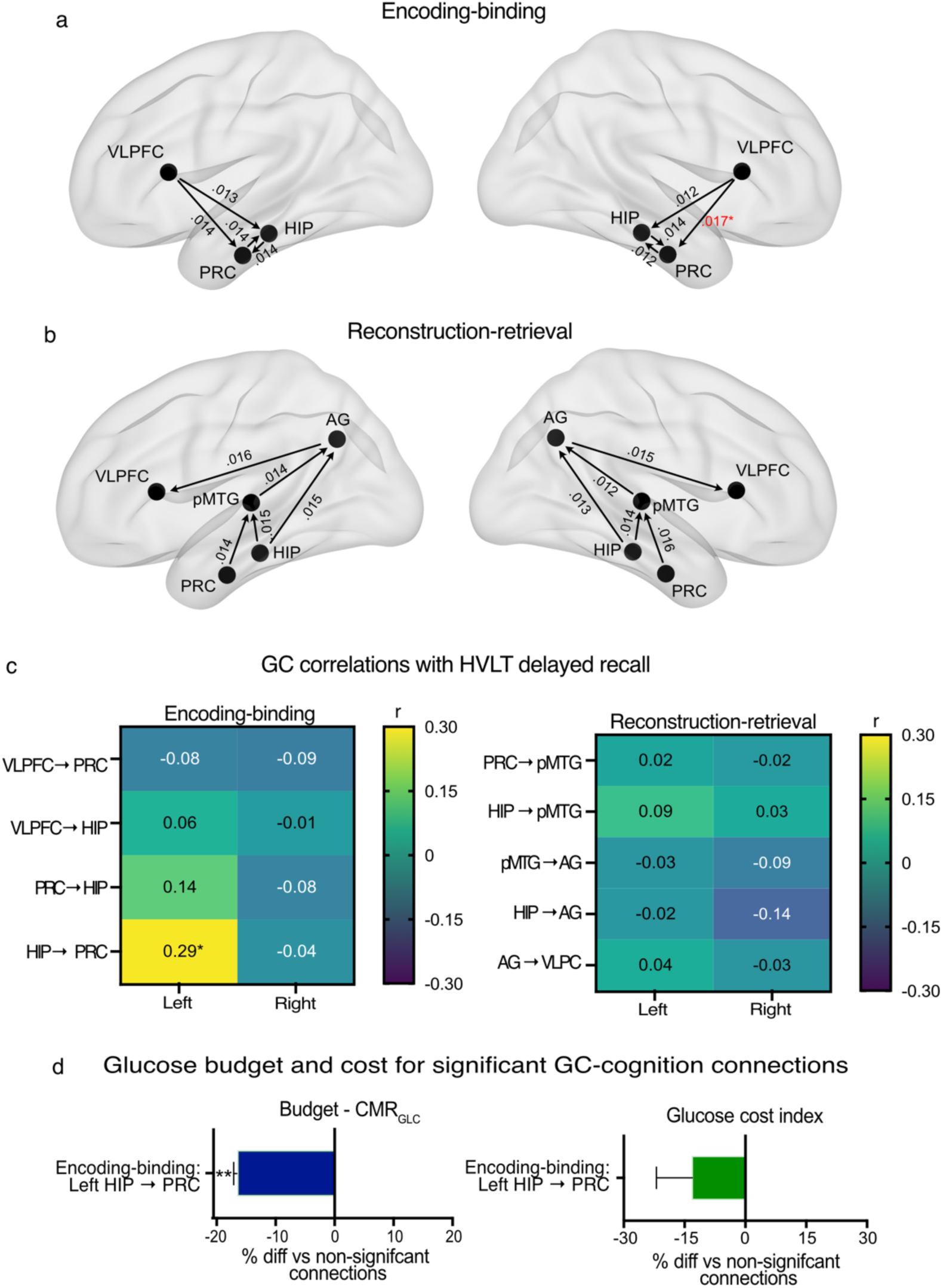
Directed metabolic connectivity in the verbal episodic memory circuit. Granger causality results in the memory circuit for encoding-binding (a) and reconstruction-retrieval (b) divisions. (c) Correlations between Granger causality values and memory performance. (d) Mean percentage difference in glucose budget (CMR_GLC_) and glucose cost index for connections with cognitive performance associations vs non-significant connections (error bars are standard error of the mean). *p < 0.05; **p-FDR < 0.05 (in panel a, p-values are compared to a permutation null model). HVLT = Hopkins Verbal Learning Test. VLPFC = ventrolateral prefrontal cortex (BA 45/47); PRC = perirhinal cortex (BA 35); HIP = hippocampus; pMTG = posterior middle temporal gyrus (BA 21/37); AG = angular gyrus (BA 39). Note that for visualisation purposes only, the location of some ROI nodes in panels a-b have been modified slightly from their original MNI coordinates to enable the pathways to be clearly displayed.

To test our hypothesis that episodic memory will depend on hippocampal-driven information transfer, we correlated GC values with Hopkins Verbal Learning Test (HVLT) delayed recall, with higher scores indicating better recall. Within the verbal episodic memory circuit, the correlation between left hippocampus-to-perirhinal cortex connectivity and HVLT delayed recall reached only nominal significance and failed to survive multiple comparison corrections (r = 0.29, p = 0.007, p-FDR = 0.062; Figure 3c). Consequently, this pathway’s association with cognition should be interpreted as moderate and indicative and requiring replication. Notably, this pathway exhibited 17% lower CMR_GLC_ compared to non-significant connections (p < 0.001; Figure 3d). Although the GCI was also 13% lower for this pathway, this difference was not significant (p = 0.147).

Together, these findings suggest that effective verbal episodic memory encoding is moderately related to the strength of the hippocampus to perirhinal cortex directed metabolic connectivity. Moreover, that pathway reflects economical metabolic support for information transfer. Rather than requiring increased energetic expenditure, stronger episodic memory performance is supported by a low-budget binding pathway, consistent with efficient hippocampal–cortical communication during memory consolidation.

### Affective regulation circuit

The affective regulation circuit was divided into frontolimbic and salience components (Andreescu et al., 2015; Coutinho et al., 2016; Germain et al., 2007; Liu et al., 2015; Liu et al., 2023; Moon et al., 2016; Okon-Singer et al., 2015; Zhong et al., 2011). The frontolimbic component included dorsolateral and ventrolateral prefrontal cortices, anterior cingulate, amygdala and caudate, with directed pathways reflecting reciprocal prefrontal–limbic regulation and cortico-striatal modulation of emotional action selection. The salience component comprised amygdala, anterior insula, and anterior cingulate, with directed influences modelling detection of emotionally salient stimuli and recruitment of regulatory control systems.

Directed metabolic influence was identified across the affective regulatory circuit, with all connections significantly different to zero at p-FDR < 0.05 (one sample *t*-tests; Figure 4a-b). The strongest pathway was in the frontolimbic circuit from the dorsolateral prefrontal cortex to caudate (mean GC = 0.019; p = 0.013, p-FDR = 0.011).

**Figure 4.**
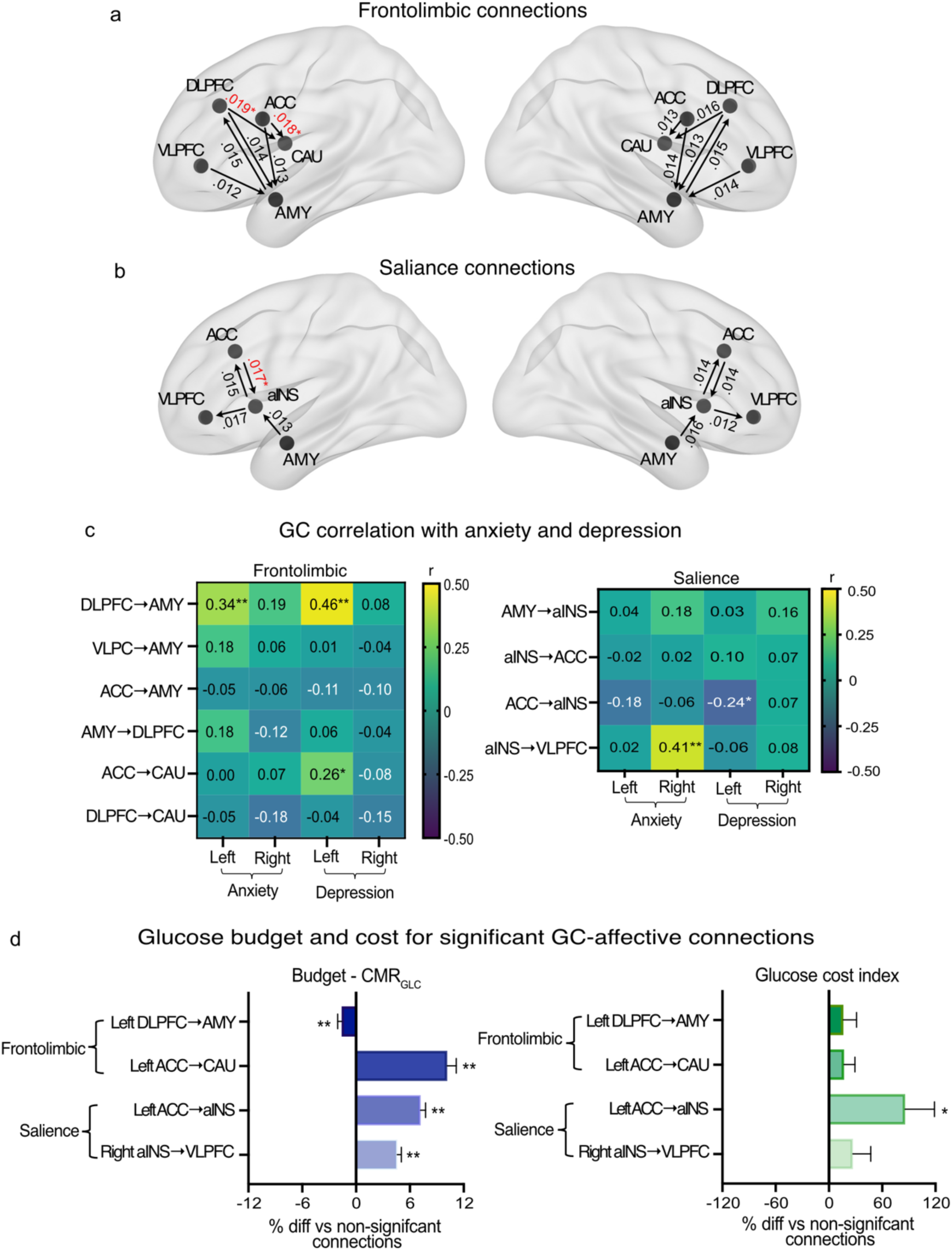
Directed metabolic connectivity in the affective regulatory circuit. Granger causality results in the affective regulatory circuit for frontolimbic (a) and salience (b) divisions. (c) Correlations between Granger causality values and Beck anxiety and depression inventory scores. (d) Mean parentage difference in glucose cost index comparing connections significantly associated with anxiety and depression vs non-significant connections (error bars are standard error of the mean). In panel a, *p < .05; **p-FDR < 0.05 compared to a permutation null model. ACC = anterior cingulate cortex (BA 24/32); AMY = amygdala; DLPFC = dorsolateral prefrontal cortex (BA9/46); aINS = anterior insula; VLPFC = ventrolateral prefrontal cortex (BA44/45/47). Note that the location of some ROI nodes in panels a-b have been modified slightly from their original MNI coordinates to enable the pathways to be clearly displayed.

We tested the hypothesis that individual differences in anxiety and depression will be related to altered directed metabolic connections in the affective regulatory circuit. We correlated the GC values with Beck anxiety and depression scores (Figure 4c). Higher scores on these inventories indicates higher levels of anxiety and depression. Based on normative cut-points for the anxiety inventory, 67 (79%) participants met the criteria for minimal anxiety, 14 (17%) mild anxiety, three (4%) moderate anxiety and one (1%) severe anxiety. For depression, 76 (89%) participants had minimal depression, four (5%) mild depression, three (4%) moderate depression and two (2%) severe depression. The Beck inventory scores indicated predominantly low levels of anxiety and depression symptoms with some heightened symptomology in a minority of participants, consistent with a largely non-clinical sample.

Stronger connectivity in the frontolimbic circuit from the left dorsolateral prefrontal cortex to the amygdala was associated with greater anxiety (r = 0.34, p = 0.002, p-FDR = 0.018) and depression (r = 0.46, p < 0.001, p-FDR < 0.001) symptoms. Stronger connectivity from the anterior cingulate to the caudate in the frontolimbic circuit was also associated with a trend towards greater depression symptoms (r = 0.26, p = 0.015, p-FDR = 0.076). In the salience circuit, stronger connectivity from the right anterior insula to the ventrolateral prefrontal cortex was associated with more anxiety symptoms (r = 0.41, p < 0.001, p-FDR = 0.002). In contrast, stronger connectivity from the left anterior cingulate to anterior insula showed a trend towards fewer depression symptoms (r = –0.24, p = 0.027, p-FDR = 0.091). As such, metabolic information flow through the frontolimbic and salience circuits was associated with poorer psychosocial outcome.

Within the frontolimbic division the behaviourally relevant connection from the left dorsolateral prefrontal cortex to amygdala demonstrated a significantly 2% lower CMR_GLC_ (p < 0.001; Figure 4d) without a corresponding difference in GCI, suggesting that effective top-down prefrontal regulation of limbic regions may occur through metabolically efficient signalling. In contrast, other frontolimbic pathways (ACC → CAU) showed a 10% higher CMR_GLC_ (p < 0.001) but no difference in GCI (+17%, p = 0.195), indicating resource-intensive, high-throughput information transfer within cortico-striatal regulatory loops. Within the salience division, the behaviourally relevant cingulate to anterior insula pathway exhibited significant increases in both CMR_GLC_ (+7%, p < 0.001) and GCI (+86%, p < 0.05), consistent with a high-cost, amplified interoceptive-regulatory signalling. In contrast, the anterior insula to ventrolateral prefrontal connection showed higher CMR_GLC_ (+5%, p < 0.001) without a significant increase in GCI (+36%, p = 0.169), suggesting greater energetic investment without proportional enhancement of efficient directed influence.

Together, these results suggest that information transfer in affective regulatory circuits is associated with anxiety and depression symptomatology. For frontolimbic pathways, stronger connectivity is associated with more anxiety and depression symptoms, reflecting affectively maladaptive information transfer with varied cost-weighted influence. By contrast, some salience connections have elevated metabolic activity without enhanced cost-weighted influence, suggesting energetically demanding information transfer associated with poor affective outcomes. Collectively, these findings indicate that individual differences in affective symptoms may emerge from a balance between metabolically efficient and costly directional signalling across frontolimbic and salience circuits, highlighting that the energetic regulation of information transfer shapes psychosocial outcomes.

### Multi-lag sensitivity

To evaluate the robustness of our directed metabolic connectivity results to alternate temporal configurations, we conducted a systematic multi-lag sensitivity analysis, recalculating directed connectivity across a range of fPET frames, including a lag of 1 frame (16s), 3 frames (48s), and 5 frames (1 min 20s). At each alternative lag, individual Granger causality metrics were extracted, re-tested against the permutation null model and correlated against the cognitive and affective measures. The detailed results are presented in Supplement 4.5, revealing a parameter-dependent trade-off in model fit: the shortest interval (Lag 1) underfit the data and failed to capture stable population-level connections, while the extended window (Lag 5) introduced systematic parameter inflation that caused significant statistical attrition in the brain-behaviour relationships. Collectively, these findings empirically validate our selection of a 2 frame lag (32 seconds) for the VAR models, demonstrating that it provides robust GC estimates while maximising sensitivity to network features and individual differences in cognitive and affective symptomatology.

### Age group differences in directed metabolic connectivity and cognition

We investigated differences in directed metabolic connectivity between older and younger adults. As the effects of age were few and of small effect size, these are reported in Supplement 4.3 and 4.4. Age moderated the strength of two GC-cognition associations. In the reactive control circuit, stronger right ventrolateral prefrontal cortex to superior parietal lobule connectivity was associated with slower stop signal reaction time in younger adults only (Z = –3.02, p-FDR = 0.023). In the frontolimbic affective regulatory network, stronger left-hemisphere anterior cingulate to caudate was associated with significantly higher depression symptoms in younger than older people (Z = 3.36, p-FDR = 0.008). Together, these results indicates that there is limited effect of age on directed metabolic connectivity and that age-specific associations with behaviour are limited to select connections in the reactive control and frontolimbic affective circuits.

## Discussion

Our results indicate that the intrinsic architecture of functional circuits is organised by directed pathways of metabolic influence. While prior work has examined average glucose metabolism (Di et al., 2019; Riedl et al., 2016; Wang et al., 2022) or undirected metabolic connectivity (Deery et al., 2026; Deery, Liang, et al., 2025; Deery, Liang, Siddiqui, et al., 2024), the present findings demonstrate that time-resolved glucodynamics can be used to measure the metabolic support for ordered, circuit-level information transfer. By identifying directed metabolic influences in functional circuits, and linking those connections to cognition and affect, our results extend metabolic imaging beyond regional activation or undirected correlations. Collectively, these results suggest that glucose metabolism does not merely index local energetic demand, but also reflects structured, behaviourally-relevant circuit dynamics, with metabolic connections supporting known pathways of information transfer between regions.

Our results reveal robust directed metabolic signalling across cognitive control, memory and affective regulatory circuits. By quantifying the energy requirements (CMR_GLC_) of the connections and by the introduction of a glucose cost index, we show that the behavioural relevance of the pathways is shaped by the efficient conversion of baseline glucose availability into functional influence. Prior work has described an energetically efficient, hub-centric network topology that balances wiring costs with information transfer capacity (van den Heuvel & Sporns).

Our glucose cost index provides a complementary pathway-level measure and shows that metabolic efficiency manifests differently across and within circuit connections. Importantly, baseline CMR_GLC_ and the glucose cost index capture distinct aspects of circuit physiology. Baseline CMR_GLC_ reflects the intrinsic energetic “set point” of a region (Jamadar et al., 2025). In contrast, the glucose cost index integrates this baseline budget with the magnitude of directed information transfer, that quantifies the energetic expenditure associated with signalling along a pathway.

Divergent patterns of CMR_GLC_ and glucose cost across circuits therefore likely reflect differences in computational and functional responses to internal and external demands. Proactive control circuits require both high baseline metabolic investment and strong directional signalling to maintain stable top-down attention. Memory circuits benefit from connections that are metabolically economic, sparse, and precisely timed (e.g., hippocampal-cortical signalling). These circuit-specific differences indicate that metabolic efficiency is not governed by a single “more is better” principle but reflects context-dependent optimisation of metabolic resources across cognitive systems.

Our finding of directed metabolic connections in both proactive and reactive control circuits aligns with and extends the dual mechanisms of control model (Braver, 2012) by the identification of distinct metabolic efficiency signatures. The dual mechanisms of control model posits that proactive and reactive circuits can operate independently, yet are often co-engaged, with their relative weighting shaped by task demands and individual differences in cognitive and affective traits (Braver, 2012; Karayanidis & Jamadar, 2014).

This weighting also reflects a resource trade-off in which proactive control has a higher attention and metabolic cost (Braver, 2012). Our findings provide circuit-level metabolic evidence for this trade-off. They suggest that metabolic support for cognitive control is optimised by selectively enabling proactive information transfer while maintaining strategically metabolically constrained reactive signalling. This trade off reflects a balance in the metabolic architecture that support sustained goal-directed control with flexible monitoring and response to internal and external demands.

Our findings suggest that working memory capacity is supported by efficient directed metabolic connections. Meta-analyses of fMRI studies have shown that working memory recruits lateralised frontoparietal networks, with posterior parietal regions supporting memory storage and dorsolateral prefrontal cortex supporting attentional control and manipulation processes (D’Esposito et al., 2000; Rottschy et al., 2012; Wager & Smith, 2003). We observed directed metabolic connectivity consistent with this multi-circuit model of working memory (Baddeley, 2003; Repovs & Baddeley, 2006). In terms of episodic memory, prior neuroimaging studies implicate hippocampal–cortical interactions in successful encoding (Eichenbaum et al., 2007; Kim, 2011; Spaniol et al., 2009; Wagner et al., 1998; Wagner et al., 2005). Consistent with this previous work, relational binding accounts of episodic memory (Ranganath et al., 2004) and the HERA framework (Habib et al., 2003; Tulving et al., 1994; Tulving et al., 1996), we found stronger left hippocampus to perirhinal cortex connectivity associated with episodic memory performance. Together, our results suggest that memory performance depends on efficient directional metabolic signalling that operates with a relatively low glucose budget. Critically, working and episodic memory circuits have select, efficient energetic pathways supporting information transfer.

Our findings advance a theoretical perspective in which individual differences in affective symptomatology emerge from the balance between efficient and costly directional metabolic signalling across frontolimbic and salience circuits. Frontolimbic metabolic connectivity is associated with more anxiety and depression symptoms, such that top-down pathways can enable more rumination and maladaptive emotional regulation, leading to poorer affect. These results are consistent with frontolimbic models of affective regulation (Banks et al., 2007; Ochsner & Gross, 2005; Phillips et al., 2003; Phillips et al., 2008) and fMRI studies (Heller et al., 2013; Johnstone et al., 2007; Liu et al., 2023; Schimmelpfennig et al., 2023). Within our metabolic efficiency framework, stronger signalling along the right anterior insula to ventrolateral prefrontal cortex pathway occurs without a high baseline glucose budget, appearing to reflect metabolically efficient translation of arousal signals into emotional regulation. Behaviourally, such metabolically efficient signalling may manifest as anticipatory worry or hypervigilant control strategies, which are both core features of anxiety (Paulus & Stein, 2006).

Our multi-lag sensitivity analysis revealed a notable pattern of brain-behaviour relationships alongside insights into model optimisation. Crucially, a lag of 2 (32 seconds) and 3 (48 second) fPET frames yielded stable directed connections against the permutation null model and resolved the highest number of meaningful cognitive and affective associations compared to shorter 1 frame (16 seconds) and longer 5 frame (1 min 20 second) lags. This higher concentration of associations suggests that individual differences in directed metabolic flow are most robustly linked to behaviour within this intermediate temporal window (∼30 seconds). Consequently, we relied on the lag 2 GC framework as an empirically optimised baseline for this fPET acquisition protocol and dataset.

Neurobiologically, these findings also suggest that the directed metabolic connections are present across complementary temporal scales, although the capacity to resolve them is tied to sampling rate. While our data suggest a ∼30-second window is optimal, we do not rule out the possibility that alternate temporal windows could yield new, unique insights. As fPET methodologies and scanner hardware continue to mature, achieving ultra-short frame durations of 1-3 seconds will make it possible to map rapid metabolic processes coupled to direct neural signalling (see (Hahn, 2025; Jamadar, Zhong, et al., 2021; Reed et al., 2023; Villien et al., 2014) for examples).

It is important to interpret the results of this study in the context of the experimental design. The cognitive assessments were conducted outside the scanner. The FDG-PET data was also acquired as participants viewed a drone-flight video. While highly effective at minimising head motion across the long acquisition window, this video content introduces sensory, attentional, and cognitive stimulation. Consequently, the directed connectivity patterns observed here should not be strictly interpreted as a representation of pure, intrinsic ‘resting-state’ metabolic architecture. Instead, they reflect a combination of baseline metabolic maintenance and some stimulus-driven network synchronisation. Nonetheless, our results suggest there is value in studying metabolic connectivity in these states. Intrinsic activity accounts for the majority of the brain’s energy budget (Raichle & Gusnard, 2002) and forms the baseline framework upon which all goal-directed and extrinsically-evoked behaviours arise (Fox et al., 2006; Mennes et al., 2011; Raichle, 2011; Smith et al., 2009; Vaishnavi et al., 2010). Importantly, resting-state fMRI connectivity often resembles task-evoked activation patterns, with up to ∼80% shared variance and cross-task similarity estimates ranging from r ≈ 0.5-0.9 (Chan et al., 2017; Cole et al., 2016). Assessing the relationship between cognition and resting metabolic connectivity may enable us to discover those pathways where intrinsic information flow supports behaviour. Nonetheless, we cannot infer that information transfer is directly associated with behavioural benefit or cost; such inference would require task-related fPET data (Hahn et al., 2020; Masharipov et al., 2024).^1^ Future studies should use task-based experimental designs to assess how information flow is directly associated with cognition, capitalising on temporal alignment between task events, behavioural responses, and directed metabolic interactions (Hahn et al., 2020; Hahn et al., 2016; Hahn et al., 2018; Jamadar, Zhong, et al., 2021).

The temporal resolution of our fPET acquisition (16s frames) and the inherently integrated nature of FDG trapping mean that GC estimations in this context capture low-frequency glucodynamic predictive dependencies rather than real-time neuronal communication or direct synaptic information flow. These directed dependencies are best understood as indexing the metabolic support for information processing in established functional circuits. In other words, it reflects the time-variant dynamics of the metabolism that supports cognitive and affective processing. Nonetheless, temporal precedence in our models must be interpreted as statistical predictive dependency rather than direct biological causality. Alternative physiological mechanisms could account for these connections, including localised variations in tracer delivery, regional differences in vascular-metabolic coupling, or unmodeled factors. Future validation using simultaneous electrophysiological recordings is required to establish the neural correlates of these low-frequency metabolic connections.

Here, we chose to take an empirical theoretically grounded ROI-based approach to examine directed metabolic connectivity. We examined connectivity for a subset of predefined paths and not the full model space (e.g., in Fig1a, the full model space would be nine pathways to/from each ROI, instead of the four tested paths). This decision was motivated to constrain the analysis to pathways with known involvement in the cognitive processes of interest (see Supplement). However it does preclude discovery of potentially important connections outside these known pathways. In addition, we examined intra-hemispheric connectivity only. Again, this decision was motivated to reduce the model space, but precludes tests of information transfer between hemispheres. While we observed some lateralised pathways, we did not conduct formal laterality comparisons. It therefore remains unclear whether these effects reflect true hemispheric specialisation, asymmetries in metabolic efficiency, or network-level inter-hemispheric interactions. Future work should explicitly test lateralised hypotheses and investigate inter-hemispheric metabolic signalling to clarify how bilateral network dynamics contribute to cognitive and affective outcomes.

Several study limitations should be considered. As noted above, the temporal resolution of fPET used here (16 second frames) limits the detection of faster metabolic connectivity supporting rapid neuronal activity and inter-regional communication. Furthermore, the use of specific ROIs assumes functional homogeneity across the ROI space, which may not accurately capture the anatomical and functional complexity of regions, particularly for the hippocampus or prefrontal cortex. Finally, the cross-sectional design precludes causal inferences about lifespan trajectories. Longitudinal and intervention studies are needed to establish whether the observed directed metabolic pathways are stable and modifiable. In conclusion, the brain’s metabolic architecture is characterised by pathways of temporally ordered metabolic influence that support information transfer across brain circuits. Functional brain circuits are defined by specific patterns of directed metabolic signalling, the strength of which is related to cognition and affect. These connections vary in the behavioural benefit they provide and the efficiency with which they use glucose to support adaptive cognition and regulate affect. An interplay exists between connectivity strength, efficient glucose utilisation and functional benefit, where trade-offs are made and circuit connections prioritised to support diverse cognitive and behavioural processes.

## Methods

### Ethical Considerations

The study received approval from the Monash University Human Ethics Research Committee. Authorisation for the use of ionizing radiation was granted by the Monash Health Principal Medical Physicist Administration, in compliance with the Australian Radiation Protection and Nuclear Safety Agency Code of Practice (2005). All participants provided informed consent prior to taking part in the research.

### Participants

For this study, we utilised the Monash *MetConn* simultaneous PET/MR dataset, which has been described in detail previously (Deery, Liang, Di Paolo, et al., 2024; Deery et al., 2026; Deery, Liang, et al., 2025; Deery, Liang, Siddiqui, et al., 2024; Deery, Moran, et al., 2025). Descriptive statistics for the sample and the younger and older groups are provided in Supplementary Table 1. Briefly, participants were recruited from the local community and comprised 86 adults. The mean age of the whole sample was 54 years (SD = 25) and the proportion of women was 51%. The mean age of the younger group was 27 years (SD = 6) and the older group was 76 years (SD = 6). The proportion of females was 54% in the younger group and 46% in the older group.

### Data acquisition

Participants completed a demographic and background questionnaire, which included the Beck Anxiety and Depression Inventories (Beck et al., 1988; Beck et al., 1961). This was followed by a cognitive test battery including the Hopkins Verbal Learning Test (HVLT) (Shapiro et al., 1999), a forward and backward digit span task (Blackburn & Benton, 1957), a task-switching paradigm (Friedman et al., 2008) and a stop-signal task (Verbruggen et al., 2008) (see Supplement 1 for details). Participants underwent a 90-minute simultaneous functional PET-MR scan in a Siemens (Erlangen) Biograph 3-Tesla molecular MR scanner. Positioned supine with their head in a 32-channel radiofrequency head coil, participants first underwent structural MRI. Non-functional T1 and T2 scans were acquired over the initial 12 minutes. The T1 3DMPRAGE scan parameters were: TA = 3.49 min, TR = 1,640ms, TE = 234ms, flip angle = 8°, field of view = 256 × 256 mm^2^, voxel size = 1.0 × 1.0 × 1.0 mm^3^, 176 slices, sagittal acquisition. For the T2 FLAIR scan: TA = 5.52 min, TR = 5,000ms, TE = 396ms, field of view = 250 × 250 mm^2^, voxel size = 0.5 × 0.5 × 1.0 mm^3^, 160 slices). List-mode PET acquisition began 13 minutes into the session (voxel size = 1.39 x 1.39 x 5.0mm^3)^. At commencement of the PET scan, half of the 260 MBq FDG tracer was administered as a bolus, with the remaining half infused continuously over 50 minutes at 36 mL/hour. This protocol balances a rapid initial increase in signal-to-noise ratio with its sustained maintenance throughout the acquisition period (Jamadar et al., 2020). During a subsequent 40-minute resting-state period, participants viewed a drone-flight video over the Hawaii Islands. Plasma radioactivity was measured from 5 mL blood samples drawn at baseline and at 10-minute intervals post-infusion.

### PET Data Preprocessing

Participant PET data were binned into 344 3D sinogram frames of 16s periods. Attenuation correction was performed using a pseudo-CT method for hybrid PET-MR scanners (Burgos et al., 2014). 3D volumes were reconstructed with the Ordinary Poisson-Ordered Subset Expectation Maximization algorithm incorporating point spread function correction (3 iterations, 21 subsets). Reconstructed DICOM slices were converted to NIFTI format, 344 × 344 × 127 (size: 1.39 × 1.39 × 2.03 mm^3^) and concatenated into a single 4D NIFTI volume. These 4D volumes underwent motion correction and partial volume effect correction (Greve et al., 2016; Jenkinson et al., 2002). We applied a 25% grey matter threshold and surface-based spatial smoothing with an 8 mm Gaussian kernel (FWHM) (Greve et al., 2014).

To minimise the impact of the FDG baseline uptake on our VAR models, the fPET time-series were denoised using the standard anatomical component based noise correction (aCompCor) framework implemented within the CONN toolbox (Whitfield-Gabrieli & Nieto-Castanon, 2012). High-resolution structural images were segmented into grey matter, white matter (WM), and cerebrospinal fluid (CSF) masks. The first five principal components capturing the variance within these non-neural tissue compartments were extracted and entered as covariates in a linear regression model to filter the grey matter signal. Temporal frequency filtering settings in CONN were defined as [0, 0.0625]. Given our sampling rate of 16 s (f_s_ = 0.0625 Hz, Nyquist f_N_ = 0.03125 Hz) the upper bound encompasses the entire discrete spectrum up to the Nyquist limit, acting as an all-pass filter that preserved the full temporal frequency range of the residuals for subsequent Granger causality analyses. As a result, the monotonic FDG accumulation trend was removed by the aCompCor nuisance regression procedure, not by temporal filtering, yielding residual signals that fluctuate around a stable baseline throughout the scan (see representative denoised time series in Supplementary Fig. S1).

The application of Granger causality to resting-state fPET data relies on an important conceptual expansion of classical tracer kinetics. While traditional static FDG-PET assumes a macroscopic steady-state where the net influx constant (Ki) is treated as a static parameter (Phelps et al., 1979; Sokoloff et al., 1977), fPET relies on the premise that localised synaptic populations undergo subtle, continuous fluctuations in glucose uptake and utilisation, even during unconstrained rest. These micro-fluctuations create minute, time-variant deflections embedded within the broader, monotonically increasing time activity curve (see Fig S1). By applying appropriate denoising, we can isolate these glucodynamic fluctuations in the timeseries from the accumulation trend and use the timeseries to measure metabolic connectivity (Deery et al., 2026; Deery, Liang, et al., 2025; Deery, Liang, Siddiqui, et al., 2024; Jamadar, Ward, et al., 2021). Therefore, in the current study, our GC analysis detects whether a localised alteration in glucose use within a source region reliably precedes and predicts a subsequent shift in a downstream target region.

### ROI metabolic rates of glucose (CMR_GLC_)

The availability of glucose and the efficiency with which it is utilised within brain circuits may help explain their contribution to cognitive and affective function(Jamadar et al., 2025). To test this possibility, we first calculated cerebral metabolic rates of glucose (CMR_GLC_) in the ROIs of the circuits using the FDG time activity curves. The FDG in the plasma samples was decay-corrected for the time between sampling and counting as the input function to Patlak models (Karjalainen et al., 2020).

## Analyses

### Granger causality

For each subject, fPET time-series were extracted as the mean metabolic signal within each ROI. Stationarity was confirmed as a prerequisite for Granger causality estimation. Directed metabolic influences were then computed separately for left– and right-hemisphere ROI pairs, with midline structures included in each hemisphere circuit (see Supplementary Table S2-5 for complete list). For the predefined source–target connections, we quantified Granger causality values using a VAR model with a lag of two frames. GC magnitude was quantified as the log ratio of the residual variances of the restricted (target’s own past) and full (including source’s past) models.

For the fPET timeseries, target (***Y_t_***) and source (***X_t_***), with a lag of p=2, the restricted model is expressed as:

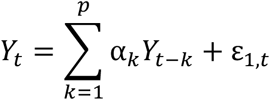

The full model is expressed as:

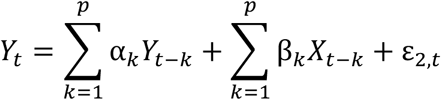

Where α*_k_* and β*_k_* represent the model coefficients, and ε_1,*t*_ and ε_2,*t*_ are the residuals. The magnitude of the directed metabolic connectivity (GC*_X_*_→F_) is calculated as the log ratio of their residual variances:

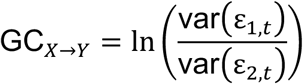

If the inclusion of the source’s past significantly reduces the prediction error variance of the target (var0ε_2,*t*_1 < var0ε_1,*t*_1), a directed predictive relationship (GC*_X_*_→F_ > 0) is established.

### Vector autoregressive (VAR) model specification and lag justification

Selection of a vector autoregressive (VAR) model depth of two fPET frames (Lag 2) was guided by a balanced optimisation of mathematical parsimony, the temporal resolution of the fPET acquisition (16 second frames), and the underlying physiological kinetics of the FDG radiotracer. To empirically validate the robustness of this model configuration and ensure that our primary brain-behaviour relationships were not artifacts of a specific temporal boundary, a comprehensive multi-lag sensitivity analysis was conducted using Lag 1 (16s), Lag 3 (48s), and Lag 5 (1 min 20s) (see Supplement 4.5 for details).

Theoretically, shorter intervals (Lag 1) are localised to more immediate glucodynamics, whereas extended intervals (Lags 5) integrate feedback loops and sustained network states over longer periods.

Ultimately, Lag 2 was retained as our primary analytical framework balancing statistical parsimony and empirical sensitivity. The inclusion of more fPET frames (e.g., Lag 5) requires the estimation of significantly more parameters, inflating raw Granger causality magnitudes by absorbing unmodeled residual variance and inducing statistical attrition among behavioural associations. Conversely, under-parameterised windows (e.g., Lag 1) underfit the data and are statistical less stable under permutation null testing. Lag 2 optimally balances these trade-offs, preserving maximum sensitivity to individual differences in cognitive and affective phenotypes while maintaining population-level architectural integrity.

### Tests of Granger causality significance

At the group level, two complementary inference approaches were employed to test the significance of the Granger causality (GC) connections. First, a one-sample right-tailed *t*-test was applied to subject-level values for each directed connection to test whether the mean GC differed significantly from zero. Second, to assess statistical significance relative to a biologically plausible null model at the subject level, the target time series was circularly shifted by a random offset and GC recomputed. This procedure was repeated 1,000 times per subject per connection, yielding a subject-specific null distribution under the hypothesis of no directed influence. Circular shifting preserves the autocorrelation structure, spectral properties, and amplitude distribution of individual ROI signals while disrupting temporal cross-dependence between regions, making it well suited to the slow dynamics of fPET (Doubovikov & Aksenov, 2025; Handwerker et al., 2012). For group-level non-parametric inference, one null GC value was randomly drawn from each retained subject’s null distribution and averaged across subjects; this process was repeated 1,000 times to construct a null distribution of the group mean under the null hypothesis. The right-tailed permutation p-value was defined as the proportion of null means greater than or equal to the observed group mean GC, with a pseudo-count of one added to both numerator and denominator to avoid zero p-values. P-values were corrected for multiple comparisons across connections using the Benjamini–Hochberg false discovery rate (FDR) procedure. Connections with FDR-adjusted p < 0.05 were considered statistically significant. Subject-level GC values were retained for subsequent cognition analyses.

### Granger causality, cognitive performance and affect

To test our hypotheses that network-specific directed metabolic connectivity will be associated with cognition and affective symptoms, Pearson correlations were calculated between test performance measures and the left– and right-hemisphere connections within each predefined circuit. Cognitive and affective measures were selected to correspond to each network’s functional domain. For the cognitive control network, category switch cost (difference reaction time between switch and non-switch trials) and stop signal reaction times were used (reversed so that higher scores indicate better performance). For the verbal episodic memory network, the HVLT delayed recognition score was used, and for working memory the sum of average longest forward and backward digit span task were used. For the affective regulation circuit, scores on the Beck anxiety and depression inventory were used. These correlations were tested at p < .05 and p-FDR < .05 for each hemisphere-circuit.

### Glucose budget and cost

Prior theoretical and empirical work has described that there is a metabolic cost to cognition that is heterogeneous across time and space in the brain (Bassett & Bullmore, 2006; Bullmore & Sporns, 2012; Deery, Liang, Siddiqui, et al., 2024; Jamadar et al., 2025; Palombit et al., 2022). We undertook exploratory analyses to quantify these costs in the metabolic circuits. To assessed whether directed connections exhibiting significant associations with cognition and affect differed in their glucose use relative to their network-specific baseline, we compared their baseline cerebral metabolic rates of glucose (CMR_GLC_). We compute a ‘glucose cost index’ (GCI) for the directed metabolic connections as the product of the Granger causality value CMR_GLC_ of its constituent ROIs, defined as:

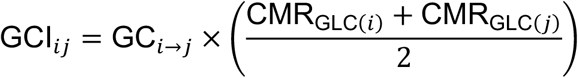

A multiplicative framework was chosen to allow us to isolate pathways that exert strong directional influence relative to their localised metabolic baseline, providing a pathway-specific index of energetic investment.

We computed a percentage difference score for each subject for both CMR_GLC_ and GCI. First, for every network and hemisphere separately, we calculated the per-subject percentage difference between each significant and all non-significant connections to expresses the metabolic cost of the significant connections. For each connection, we then tested whether the mean percentage difference between significant and non-significant connections across subjects differed from zero using a one-sample *t*-test, thresholded at α=0.05 (uncorrected for these exploratory analyses). This approach allowed us to evaluate whether significant connections consistently have a higher or lower baseline glucose budget and consume more or less metabolic resources than non-significant connections within their respective network.

### Outlier detection

Subjects exceeding ±3 standard deviations on a Granger causality or CMR_GLC_ were excluded on a pairwise-basis from the correlation and GCI analyses, with the resulting n-sizes reported in the tables of results.

## Data and Code Availability Statement

The dataset and code used for the study are available at https://osf.io/gbfcu/overview.

## Author Contributions

SDJ and GFE conceived the project. HD and SDJ, CM and GFE designed and developed the manuscript. HD and EL analysed the data. HD wrote the manuscript, SDJ, GFE and CM reviewed/edited the manuscript. All authors read and approved the final manuscript.

## Acknowledgements

This work was supported by Australian Research Council (ARC) Discovery Project DP25010302 and ARC Fellowship FT250100206.

The authors acknowledge the facilities and scientific and technical assistance of the National Imaging Facility (NIF), a National Collaborative Research Infrastructure Strategy (NCRIS) capability at Monash Biomedical Imaging (MBI), a Technology Research Platform at Monash University.

## Footnotes

1 While the current study is the first to examine directed information transfer in the metabolic network, it does bear conceptual similarity to some previous work using functional PET. For example, Hahn et al. (2020) used ‘metabolic connectivity mapping’ (Riedl et al., 2014) to infer directionality of information transfer during a task (Tetris game). However, importantly, the connections of interest were derived from fMRI data, with the fPET data used to infer directionality of fMRI connectivity across the network. Thus, the MCM approach uses PET/fPET data to understand information flow across the functional fMRI network of the brain, whereas here we used fPET to understand information flow across the metabolic network of the brain.

