## Supplement for "Directed information flow across the metabolic network of the human brain"

~ Supplementary Methods and Results ~

**1. SUPPLEMENTARY METHODS 2**
1.1 Participants 2
1.2 Cognitive tests 2
1.3 Data acquisition 3

**2. PARTICIPANT DEMOGRAPHICS 4**

**3. CIRCUIT DEFINITION 5**
3.1 Cognitive control circuit 5
3.2 Working memory circuit 7
3.3 Verbal episodic memory circuit 9
3.4 Affective regulatory circuit 11

**4. SUPPLEMENTARY RESULTS 13**
4.1 Signal denoising and characterisation of fPET timeseries 13
4.2 Can activity in the proactive circuit ameliorate interference-related activity in the reactive circuit? 14
4.3 Age group differences in cognition and directed metabolic connectivity 15
4.4 Age group differences in the association between directed metabolic connectivity and cognition 15
4.5 Sensitivity analysis of Granger Causality lag parameter 24

**SUPPLEMENTARY REFERENCES 27**

### 1. Supplementary methods

#### *1.1 Participants*

Local advertising was used to recruit 90 participants from the general community. A screening interview was conducted to ensure that participants had the capacity to provide informed consent. Participants were also screened to ensure that they did not have a diagnosis of diabetes, neurological or psychiatric illness, claustrophobia or non-MR compatible implants. They were also excluded if they had received a clinical or research PET scan in the past 12 months. Women were screened for current or suspected pregnancy. Participants received a $100 voucher for participating in the study. Four participants were excluded due to excessive head motion (N=2) or incomplete PET scan or image reconstruction (N=2).

#### *1.2 Cognitive tests*

**Verbal Episodic Memory - Hopkins Verbal Learning Test (HVLT).** The HVLT is a three-trial list learning and free recall task. The learning trials comprised 12 words, four words from each of three semantic categories (Shapiro et al., 1999). Approximately 20–25 minutes after the learning trails, participants completed delayed recall and recognition trials. The delayed recall required free recall of any of the 12 words. The recognition trial comprised 24 words, including the 12 target words and 12 false-positives, six semantically related, and six semantically unrelated. Delayed recall was calculated as the total words recalled.

**Working Memory - Digit Span.** A measure of verbal short term and working memory used in two formats: Forward and backward digit span (Blackburn & Benton, 1957). Participants were presented with a series of digits, and are asked to repeat them in either the order presented (forward span) or in reverse order (backwards span). After two consecutive failures of the same length, the test was stopped. Scores were derived as the sum of the longest correct series for forward and backward recall.

**Proactive Cognitive Control - Task Switching.** For task switching, a computerised test was used in which participants were presented with a word and had to perform a categorisation task. The categorisation task was dependant on two cues that appeared on screen across the trials. One cue was a heart symbol, for which participants were asked to categorise the word presented via a key press as either a LIVING or a NON-LIVING object. If the cue was an arrow-cross, participants were asked to categorise the word as either BIGGER or SMALLER than a basketball. The cue was randomised across trials. Half the trials were switch trials and half were non-switch trials. Half the switch and non-switch trials was congruent in the key presses for either task, half was incongruent. The task switching measure was switch cost, calculated as the difference reaction time between switch and non-switch trials (Friedman et al., 2008).

**Reactive Cognitive Control - Stop Signal.** The stop signal trial was a computer-based test (Verbruggen et al., 2008). Participants were required to press the left response key if an arrow on screen pointed left and the right response key if the arrow pointed right. If a signal beep sounded, participants were instructed stop their response. The delay between presentation of an arrow and signal beep started at 250ms and was altered up or down by 50ms based on performance. The delay increased up to 1150ms if the previous stop signal trial was successful and decreased down to 50ms if the previous stop signal trial was unsuccessful. The stimulus onset asynchrony between the onset of a fixation circles at the start of each trial was 2000ms. Reaction time in the stop signal trials was recorded.

#### 1.3 Data acquisition

Participants underwent a 90-minute simultaneous MR-PET scan in a Siemens (Erlangen) Biograph 3-Tesla molecular MR scanner. Participants were instructed to consume a high-protein and low-sugar diet for 24 hours and to fast completely for six hours prior to the scan. They were also asked to consume 2–6 glasses of water in the six hours before the scan. Upon arrival, participants were cannulated in the vein in each forearm and a baseline blood sample of 10ml was taken. At the start of the scan, half of the 260 MBq FDG tracer was administered as a bolus to provide a strong PET signal. The remaining 130 MBq of the tracer dose was infused over 50 minutes at a rate of 36ml/hour. This combined bolus plus constant infusion protocol provides a good balance between a rapid increase in signal-to-noise ratio at the beginning of the scan, and maintenance of signal-to-noise ratio over the length of the scan (Jamadar et al., 2020).

Participants were positioned supine in the scanner bore and instructed to lie as still as possible. Their head was placed in a 32-channel radiofrequency head coil. The scan sequence commenced with non-functional T1 and T2 MRI scans in the first 12 minutes to image the anatomical grey and white matter structures, respectively. The T1 3DMPRAGE scan parameters were: TA = 3.49 min, TR = 1,640ms, TE = 234ms, flip angle = 8°, field of view = 256 × 256 $mm^2$, voxel size = 1.0 × 1.0 × 1.0 $mm^3$, 176 slices, sagittal acquisition. The T2 FLAIR parameters were: TA = 5.52 min, TR = 5,000ms, TE = 396ms, field of view = 250 × 250 $mm^2$, voxel size = .5 × .5 × 1 $mm^3$, 160 slices). Thirteen minutes into the scan, list-mode PET (voxel size = 1.39 x 1.39 x 5.0$mm^3$) and T2* EPI BOLD-fMRI (TA = 40 minutes; TR = 1,000ms, TE = 39ms, FOV = 210 mm2, 2.4 × 2.4 × 2.4 $mm^3$ voxels, 64 slices, ascending axial acquisition) sequences were initiated. A 40-minute resting-state scan was undertaken while participants watched a movie of a drone flying over the Hawaii Islands. Pseudo-continuous arterial spin labelling (pc-ASL) and diffusion-weighted imaging (DWI) was also acquired at the end of the scan but are not reported here.

Participants' plasma radioactivity levels were measured across the scan. Beginning at 10-minutes post infusion, 5ml blood samples were taken at 10 minute intervals from the right forearm. The blood sample were directly placed in a centrifuge and spun for 5 minutes at 2,000 rpm (RCF ~ 515g). 1,000-μL plasma was pipetted to a counting tube and placed in a well counter for four minutes. For each sample, the count start time, total number of counts, and counts per minute were documented.

### 2. Participant demographics

**Table S1.** Descriptive statistics for the sample and younger and older groups.

| | Whole sample (N = 86) | | Younger (N = 40) | | Older (N = 46) | | Younger vs Older[1] |
|---|---|---|---|---|---|---|---|
| | Mean | SD | Mean | SD | Mean | SD | p-value |
| Age | 53.5 | 24.8 | 27.9 | 6.2 | 75.8 | 6.0 | < 0.001 |
| Sex (% female) | 51% | | 54% | | 46% | | 0.662 |
| Fasting blood glucose (mmol/L) | 5.0 | 0.5 | 4.8 | 0.4 | 5.2 | 0.6 | < 0.001 |
| Systolic blood pressure (mmHg) | 81.6 | 12.8 | 78.5 | 13.1 | 84.2 | 12.0 | < 0.001 |
| Diastolic blood pressure (mmHg) | 78.4 | 15.4 | 82.5 | 16.9 | 74.8 | 13.1 | 0.036 |
| Body Mass Index (kg/m$^2$) | 25.0 | 4.1 | 24.1 | 4.6 | 25.7 | 3.6 | 0.075 |
| HVLT: delayed recall | 8.2 | 2.7 | 9.3 | 2.5 | 7.3 | 2.6 | < 0.001 |
| Digit span: longest span recall | 12.1 | 2.2 | 12.3 | 2.7 | 11.9 | 1.8 | 0.449 |
| Category switch: switch cost (sec) | 0.38 | 0.37 | 0.28 | 0.23 | 0.48 | 0.43 | 0.011 |
| Stop signal: RT (sec) | 0.57 | 0.13 | 0.53 | 0.13 | 0.60 | 0.12 | < 0.001 |
| Beck anxiety inventory (BAI)[2] | 5.5 | 6.7 | 6.8 | 8.8 | 4.3 | 3.8 | 0.088 |
| Beck depression inventory (BDI)[2] | 6.3 | 6.9 | 8.6 | 8.4 | 4.2 | 4.5 | 0.003 |

[1]Younger vs older comparisons are based on independent sample *t*-tests (two tailed), except sex which uses Chi-squared.
[2]Beck Inventory normative cut points:
For the BAI, 0-7 = minimal anxiety; 8-15 = mild anxiety, 16-25 = moderate anxiety; 26-63 = severe anxiety.
For the BDI, 0-13 = minimal depression; 14-19= mild depression, 20-28 = moderate depression; 29-63 = severe depression.

Based on the normative cut-off points from the Beck inventories (Beck et al., 1988; Beck et al., 1961), our sample has predominantly minor levels of anxiety and depression symptoms with some heightened symptomology in a minority of participants, consistent with a largely non-clinical sample. For anxiety, 67 (79%) participants met the criteria for minimal anxiety, 14 (17%) mild anxiety, three (4%) moderate anxiety and one (1%) severe anxiety). For depression, 76 (89%) had minimal depression, four (5%) mild depression, three (4%) moderate depression and two (2%) severe depression.

### 3. Circuit definition

In this section, the directed metabolic connections between regions in circuits supporting cognitive control, working and verbal episodic memory and affective regulation are described. The ROI coordinates and associated Brodmann areas are provided, together with a description of the connections tested using Granger causality analysis and their function in the circuit and reported in the main manuscript. In each section, the neuroscience and neuroimaging literature reporting these circuit connections is briefly reviewed.

#### *3.1 Cognitive control circuit*

**Table S2.** ROIs and directed metabolic connections tested in Granger causality analyses of the cognitive control circuit.

| ROI / Connection | MNI Coordinates (x, y, z) | Brodmann Area(s) | Hem | Radius (mm) | Primary Functional Role |
|---|---|---|---|---|---|
| Anterior Cingulate Cortex (ACC) | 0, 24, 32 | BA 32 | Mid | 10 | Conflict monitoring; performance evaluation |
| Dorsolateral Prefrontal Cortex (DLPFC) | ±42, 24, 32 | BA 9 / 46 | L/R | 10 | Goal maintenance; top–down biasing |
| Ventrolateral Prefrontal Cortex (VLPFC) | ±48, 24, -4 | BA 44 / 45 / 47 | L/R | 10 | Response inhibition; rule implementation |
| Inferior Frontal Junction (IFJ) | ±44, 6, 32 | BA 44 / 6 | L/R | 10 | Task switching; rule updating |
| Superior Parietal Lobule (SPL) | ±24, -64, 48 | BA 7 | L/R | 10 | Attentional reconfiguration; spatial updating |
| Anterior Insula (aINS) | ±34, 20, -4 | BA 13 | L/R | 10 | Salience detection; control recruitment |
| Caudate | ±12, 12, 8 | — | L/R | 8 | Action selection; cortico-striatal gating |

| Division | Connection | Function |
|---|---|---|
| Proactive | ACC → CAU | Monitoring signals → striatal action gating |
| Proactive | ACC → DLPFC | Conflict monitoring → reinforcement of task goals |
| Proactive | DLPFC → ACC | Top–down modulation of monitoring systems |
| Proactive | DLPFC → CAU | Executive signals → action selection gating |
| Reactive | aINS → ACC | Salience detection → recruitment of control resources |
| Reactive | aINS → VLPFC | Salience processing → response regulation |
| Reactive | VLPFC → IFJ | Rule implementation → switching initiation |
| Reactive | IFJ → SPL | Task-set update → parietal attentional broadcasting |
| Reactive | VLPFC → SPL | Inhibitory control → attentional remapping |

Cognitive control circuits were divided into proactive and reactive components in accordance with the Dual Mechanisms of Control (DMC) framework (Braver, 2012; Braver et al., 2003). Proactive control reflects the sustained, anticipatory maintenance of goal-relevant information to bias downstream processing prior to interference (Braver, 2012). In the present model, proactive control was operationalised within a circuit centred on dorsolateral prefrontal cortex (DLPFC), anterior cingulate cortex (ACC), and caudate nucleus, consistent with evidence for cortico-striatal gating mechanisms supporting goal maintenance (Miller & Cohen, 2001). Directed influences among DLPFC → ACC, DLPFC → caudate, ACC → DLPFC, and ACC → caudate were defined as proactive pathways, reflecting top–down goal maintenance and performance monitoring signals that gate action selection via cortico-striatal loops(Cohen et al., 1990).

Reactive control reflects transient, stimulus-triggered recruitment of control following detection of salient or high-interference events (Braver, 2012). In the current circuit, reactive control was modelled within a network comprising anterior insula (aINS), ventrolateral prefrontal cortex (VLPFC), inferior frontal junction (IFJ), and superior parietal lobule (SPL), consistent with salience-network driven control recruitment and frontoparietal reconfiguration mechanisms (Aron et al., 2004; Menon & Uddin, 2010; Seeley et al., 2007). Directed pathways including aINS → ACC, aINS → VLPFC, VLPFC → IFJ, IFJ → SPL, and VLPFC → SPL were classified as reactive, consistent with models in which salience detection recruits ventrolateral control regions that initiate rule updating and broadcast attentional reconfiguration signals to the parietal cortex (Aron et al., 2004).

#### *3.2 Working memory circuit*

**Table S3.** ROIs and directed metabolic connections tested in Granger causality analyses of the working memory circuit.

| ROI / Connection | MNI Coordinates (x, y, z) | Brodmann Area(s) | Hem | Radius (mm) | Primary Functional Role |
|---|---|---|---|---|---|
| Ventrolateral Prefrontal Cortex (VLPFC) | -50, 20, 10 | BA 45 / 47 | L | 10 | Articulatory rehearsal; strategic control of subvocal articulation |
| Inferior Parietal Lobule (IPL) | -45, -45, 45 | — | L | 10 | Phonological storage; maintenance of items in working memory |
| Intraparietal Sulcus (IPS) | -34, -48, 42 | — | L | 10 | Attentional focus within working memory; serial order scanning |
| Dorsolateral Prefrontal Cortex (DLPFC) | -42, 24, 32 | BA 9 / 46 | L | 10 | Executive manipulation (backward span); goal maintenance |
| Medial Frontal Cortex (mPFC) | 0, 14, 54 | BA 6 / 32 | Mid | 10 | Top-down control; attentional regulation and monitoring |

| Division | Connection | Function |
|---|---|---|
| Attention / Executive | IPS → DLPFC | Attended item delivered to executive for manipulation |
| Attention / Executive | DLPFC → IPS | Attentional shift command (core mechanism of backward span) |
| Attention / Executive | DLPFC → VLPFC | Strategic control of rehearsal (chunking, pacing) |
| Attention / Executive | mPFC → DLPFC | Guides executive manipulation and monitoring |
| Attention / Executive | mPFC → IPL | Modulates storage/attention network |
| Encoding / Maintenance | VLPFC → IPL | Rehearsal refreshes phonological store (forward span) |
| Encoding / Maintenance | IPL → VLPFC | Store status feedback regulates rehearsal rate |
| Encoding / Maintenance | IPL → IPS | Contents of store brought into focal awareness for inspection |

The working memory circuit was divided into attention-control and storage-maintenance components, reflecting functional divisions in the cognitive neuroscience literature (Hartley & Speer, 2000; Liu et al., 2025). The attention-control network, comprising dorsolateral prefrontal cortex (DLPFC), intraparietal sulcus (IPS), and medial frontal cortex (mPFC), subserves executive operations including top-down attentional control, task management, and the selective gating of information into working memory (Vartanian et al., 2022; Wang et al., 2025; Zhang et al., 2024). In contrast, the storage-maintenance network, consisting of ventrolateral prefrontal cortex (VLPFC) and inferior parietal lobule (IPL), supports the active maintenance, rehearsal, and temporary storage of information through reciprocal interactions with control regions and posterior representational areas (D'Esposito et al., 2000; Postle, 2006; Wager & Smith, 2003). This division was reflected in specific pathways: VLPFC → IPL drives articulatory rehearsal to refresh the phonological store, while IPL → VLPFC provides feedback regarding store status to regulate rehearsal pace. IPL → IPS channels maintain items into focal attention, which are then delivered to DLPFC via IPS → DLPFC for executive manipulation. The DLPFC orchestrates attention and rehearsal through DLPFC → IPS and DLPFC → VLPFC, respectively, implementing attentional shifts and strategic control, such as chunking or pacing. Medial frontal regions exert top-down modulation via mPFC → DLPFC and mPFC → IPL, guiding executive operations and modulating storage and attention networks.

Neuroimaging evidence supports this mechanistic architecture: dissociable activation patterns are observed in control versus storage regions (Rottschy et al., 2012). Furthermore, effective connectivity between prefrontal and parietal nodes scales with working memory demands (Liu et al., 2025), and age-related differences in frontoparietal engagement affect the dynamic allocation of attention and maintenance resources (Mitchell et al., 2016). By modelling directed influences in these circuits using Granger causality, it is possible to quantify how executive control regions dynamically shape storage processes, a core mechanism underlying flexible working memory (Kruijne et al., 2021; Postle, 2006).

#### *3.3 Verbal episodic memory circuit*

**Table S4.** ROIs and directed metabolic connections tested in Granger causality analyses of the verbal episodic memory circuit.

| ROI / Connection | MNI Coordinates (x, y, z) | Brodmann Area(s) | Hem | Radius (mm) | Primary Functional Role |
|---|---|---|---|---|---|
| Ventrolateral Prefrontal Cortex (VLPFC) | -50, 20, 10 | BA 45 / 47 | L/R | 10 | Strategic encoding; controlled retrieval cueing |
| Perirhinal Cortex (PRC) | -28, -14, -28 | BA 35 (±28 overlap) | L/R | 8 | Item familiarity; item–context binding |
| Hippocampus (HIP) | -24, -22, -18 | NA | L/R | 8 | Relational binding; episodic sequence encoding |
| Posterior Middle Temporal Gyrus (pMTG) | -58, -23, 5 | BA 21 / 37 | L/R | 10 | Lexical–semantic representation; memory trace reactivation |
| Angular Gyrus (AG) | -44, -62, 34 | BA 39 | L/R | 10 | Episodic reconstruction; cortical memory buffer |
| **Division** | **Connection** | | **Function** | | |
| Encoding–Binding | VLPFC → PRC | | Strategic encoding drives item-level processing in MTL | | |
| Encoding–Binding | VLPFC → HIP | | Top–down modulation of relational binding | | |
| Encoding–Binding | PRC → HIP | | Item representations integrated into episodic bindings | | |
| Encoding–Binding | HIP → PRC | | Feedback consolidation and memory strengthening | | |
| Reconstruction–Retrieval | PRC → pMTG | | Item-level activation triggers cortical lexical reinstatement | | |
| Reconstruction–Retrieval | HIP → pMTG | | Relational memory reinstates semantic trace | | |
| Reconstruction–Retrieval | pMTG → AG | | Semantic trace integrated into episodic buffer | | |
| Reconstruction–Retrieval | HIP → AG | | Episodic reconstruction and scene integration | | |
| Reconstruction–Retrieval | AG → VLPFC | | Retrieval completion signal to strategic control systems | | |

Verbal episodic memory circuits can be divided into encoding–binding and reconstruction–retrieval components, reflecting established functional systems in the cognitive neuroscience literature (Thommesen et al., 2024). The encoding–binding network includes the ventrolateral prefrontal cortex (VLPFC; BA 45/47), perirhinal cortex (PRC; BA 35), and hippocampus (HIP), which collectively subserve the initial registration, relational binding, and consolidation of verbal information into durable episodic representations. The VLPFC supports strategic encoding, providing top-down modulation that guides item-level processing in the medial temporal lobe (PRC) and facilitates relational binding in the hippocampus (Buckner et al., 1999; Otten et al., 2001; Wagner et al., 1998). Reciprocal interactions between the hippocampus and PRC further strengthen item–context associations, consolidating episodic traces for later retrieval (Davachi & Wagner, 2002; Eichenbaum et al., 2007; Ranganath et al., 2004; Rudner et al., 2007).

The reconstruction–retrieval network comprises the posterior middle temporal gyrus (pMTG; BA 21/37), angular gyrus (AG; BA 39), hippocampus, PRC, and VLPFC. Within this network, the PRC and hippocampus initiate item- and relational-level activation that reinstates semantic and contextual memory traces in pMTG. These representations are then integrated into the episodic buffer in AG, supporting coherent reconstruction of previously encoded information (Baddeley, 2000; Repovs & Baddeley, 2006)*.* The AG communicates with VLPFC to signal retrieval completion, enabling strategic control over memory search and elaboration (Dobbins et al., 2002; Hutchinson et al., 2009; Wagner et al., 2005). This circuit aligns with the HERA model (Habib et al., 2003; Tulving et al., 1994; Tulving et al., 1996), which posits left prefrontal regions as preferentially involved in episodic encoding and right prefrontal regions in episodic retrieval. Additionally, dual-process models of recollection and familiarity further distinguish item-based processing in PRC from relational/episodic reconstruction in HIP and AG (Rugg & Yonelinas, 2003).

This ROI- and connection-based framework reflects the directional flow of information across functionally specialised nodes: VLPFC → PRC → HIP for encoding, HIP → pMTG → AG for reconstruction, and AG → VLPFC for retrieval monitoring. By modelling directed influences in these circuits using Granger causality, it is possible to test if metabolically-driven strategic encoding and relational binding drive episodic reconstruction, providing mechanistic insight into the formation and recovery of verbal episodic memories (Kim, 2011; Spaniol et al., 2009).

#### 3.4 Affective regulatory circuit

**Table S5.** ROIs and directed metabolic connections tested in Granger causality analyses of the affective regulatory circuit.

| ROI / Connection | MNI Coordinates (x, y, z) | Brodmann Area(s) | Hem | Radius (mm) | Primary Functional Role |
|---|---|---|---|---|---|
| Amygdala | ±24, -4, -18 | Subcortical | L/R | 6 | Emotional salience; threat processing |
| Dorsolateral Prefrontal Cortex (DLPFC) | ±42, 24, 32 | BA 9 / BA 46 | L/R | 10 | Top-down emotional/cognitive control |
| Ventrolateral Prefrontal Cortex (VLPFC) | ±48, 24, -4 | BA 44 / BA 45 / BA 47 | L/R | 10 | Top-down emotional regulation; reinterpretation, suppression, labelling |
| Anterior Cingulate Cortex (ACC) | 0, 24, 32 | BA 24 / BA 32 | Mid | 10 | Conflict monitoring; error detection |
| Anterior Insula (aINS) | ±34, 20, -4 | Subcortical | L/R | 10 | Interoception; salience detection |
| **Division** | **Connection** | **Function** | | | |
| Frontolimbic | DLPFC → AMY | Cognitive → emotional regulation | | | |
| Frontolimbic | VLPFC → AMY | Strategic top-down modulation of amygdala response | | | |
| Frontolimbic | ACC → AMY | Conflict monitoring → emotional adjustment | | | |
| Frontolimbic | AMY → DLPFC | Bottom-up emotional/threat signal → cognitive appraisal | | | |
| Frontolimbic | ACC → CAU | Conflict detection → striatal action gating | | | |
| Frontolimbic | DLPFC → CAU | Executive control → reinforcement/action selection | | | |
| Salience | AMY → aINS | Emotional salience → salience network recruitment | | | |
| Salience | aINS → ACC | Salience/interoception → conflict monitoring | | | |
| Salience | ACC → aINS | Conflict monitoring → salience network modulation | | | |
| Salience | aINS → VLPFC | Salience/interoception → inhibitory control | | | |

Affective regulation circuits can be divided into frontolimbic and salience network components, reflecting distinct functional roles in emotion processing and regulation (Phillips et al., 2003; Schimmelpfennig et al., 2023). The frontolimbic network - encompassing dorsolateral prefrontal cortex (DLPFC), ventromedial prefrontal cortex (vmPFC), orbitofrontal cortex (OFC), anterior cingulate cortex (ACC), amygdala, and hippocampus - supports top-down modulation of emotional responses and the integration of emotional information with decision-making processes (Ochsner & Gross, 2005; Phillips et al., 2008). Within this network, effective emotion regulation is associated with increased prefrontal activity and concurrent modulation of limbic regions, with the strength of frontolimbic connectivity predicting successful down-regulation of negative affect (Banks et al., 2007). A dorsal-ventral dissociation further refines this system: a ventral subsystem - including amygdala, anterior insula (aINS), ventral striatum, ventral ACC, and OFC - processes emotionally salient information, while a dorsal subsystem - including DLPFC and ACC (ACC) - supports voluntary, cognitive regulation of emotion (Phillips et al., 2003).

The salience network, anchored in aINS and ACC with key nodes in amygdala and ventral striatum, functions as a dynamic hub that detects behaviourally relevant stimuli and coordinates large-scale network interactions (Seeley et al., 2007; Uddin, 2015). The salience network integrates autonomic, visceral, and emotional information to guide attentional and behavioural responses and acts as a switching mechanism between the default mode network (DMN) and central executive network (CEN) depending on task demands and stimulus salience (Menon, 2011; Sridharan et al., 2008). While the frontolimbic network implements reciprocal prefrontal-limbic interactions for regulatory control, the salience circuit initiates directed influences that modulate these regulatory circuits.

By explicitly modelling directed connections in these networks, Granger causality analysis can quantify temporal precedence in metabolic signals, revealing how frontolimbic and salience network dynamics regulatory control (Schimmelpfennig et al., 2023). This approach provides a mechanistic framework for understanding affective regulation and the circuit-specific patterns of dysregulation observed across psychiatric disorders: fronto-limbic disruptions characterise mood and anxiety disorders (Johnstone et al., 2007), whereas salience abnormalities, either hyper- or hypoactivity, are implicated in a broader spectrum of conditions including psychosis, chronic pain, and post-traumatic stress disorder (Johnstone et al., 2007; Phillips et al., 2008; Schimmelpfennig et al., 2023).

### 4. Supplementary Results

#### *4.1 Signal denoising and characterisation of fPET timeseries*

To ensure the extraction of highly reliable, neural-specific fPET timeseries and minimize the confounding impact of absolute FDG baseline uptake, a rigorous denoising and filtering pipeline was applied to the regional grey matter data. High-resolution structural scans were segmented into tissue compartments to extract the first five principal components from the white matter and cerebrospinal fluid masks via the standard aCompCor framework; these components were then regressed out of the grey matter signal alongside movement parameters to isolate localised variance that reflect ongoing synaptic glucose use. This filtering strategy successfully decouples time-variant glucodynamics from the macroscopic, monotonically increasing tracer accumulation curve. Representative examples of these timeseries used for Granger causality modelling are displayed in Figure S1.

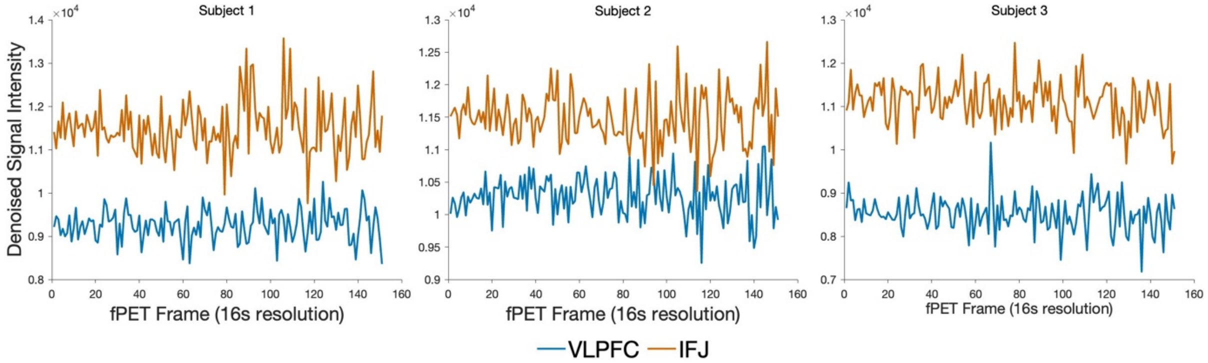


**Figure S1. Denoised fPET timeseries.** Timeseries for two ROIs from a single subject after the denoising and filtering pipeline was applied to decouple the time-variant glucose signal from the macroscopic, monotonically increasing tracer accumulation curve.

#### *4.2 Can activity in the proactive circuit ameliorate interference-related activity in the reactive circuit?*

In the main manuscript, we found that stronger right VLPFC to IFJ connectivity was associated with *slower* stop-signal reaction time. Interestingly, these results indicate that directed information transfer between nodes of the reactive control circuit is not simply associated with improved performance. Rather, in some instances, directed metabolic connectivity is associated with improved performance (faster response inhibition), and in others, *worse* performance (slower response inhibition). This indicates that in some instances, information flow in metabolic circuits may actually interfere with performance. Therefore, we were interested in understanding whether directed metabolic connectivity in one part of the circuit might reduce the negative influence of these interference effects in other parts of the circuit. In other words, can goal-directed processing, as implemented via the proactive circuit, compensate or ameliorate interference-related connections in the reactive control network? We undertook exploratory *post hoc* analyses to test the possibility that the dorsolateral prefrontal cortex to anterior cingulate connection (positive-proactive control) and anterior insula to anterior cingulate connection (positive-reactive control) is moderated by ventrolateral prefrontal cortex to inferior frontal junction connectivity (negative-reactive control). We used regression analyses predicting cognitive performance from each GC connection and their interaction, with the interaction term testing the moderating effect.

We found no evidence that the pathways moderate each other's behavioural effects. Specifically, within the reactive control network we tested the model stop signal RT = aINS → ACC + VLPFC → IFJ + aINS → ACC x VLPFC → IFJ. The regression model was significant ($F = 5.3$, $p = 0.002$), although the moderating interaction effect was not significant ($\beta = 0.154$, $p = 0.373$). Similarly, for the regression model testing moderation across proactive-reactive control (switch cost = DLPFC → ACC + VLPFC → IFJ + DLPFC → ACC x VLPFC → IFJ), the model was significant ($F = 3.5$, $p = 0.020$), however, the moderating term was not ($\beta = 0.242$, $p = 0.255$). These results suggest that, regardless of positive or negative associations with cognition, the significant GC connections are additive and that the significant connections do not moderate each other's effect on cognitive performance. These findings also suggest that information flow in proactive pathways is associated with better performance, whereas reactive pathway engagement can differentially affect performance, with negative pathways appearing to reflect an effort-dependent stopping process. Cognitive control outcomes therefore appear to reflect the balance of parallel circuit influences rather than compensatory influences between pathways.

In the cognitive control circuit, we found that information flow between regions in one part of the network that are positively associated with cognitive performance appear statistically independent of those in another part of the network that are negatively associated with performance. Given the finding of similar opposing associations with working memory performance, we next undertook exploratory post-hoc analyses to test for moderating effects in those circuits. Using the regression approach, we tested the model: digit span performance = IPL→IPS + DLPFC→IPS + IPL→IPS x DLPFC→IPS. The regression model was significant ($F = 4.3$, $p = 0.007$) but the moderating effect of the connections was not significant ($\beta = -0.113$, $p = 0.567$).

#### *4.3 Age group differences in cognition and directed metabolic connectivity*

In our final series of analyses, we investigated age group differences in directed metabolic connectivity and the association between connectivity and cognition. We used independent sample *t*-tests to compare younger (N = 40; mean age 27.9 years; range 20-42) and older (N = 46; mean 75.8; 60-89) adults in our sample (see Table S1 for demographics). First we compared the groups on the cognitive and affective measures. We then compared the groups on the GC values in the circuits.

For the cognitive performance measures, older adults had worse verbal learning and memory (HVLT) than younger adults, and worse switch and stop signal performance (all p-FDR < 0.05; see Table S8). The age group performance difference on the digit span task was not statistically significant (p-FDR = 0.449). Younger adults had higher anxiety scores than older adults (p-FDR = 0.009) and tended to have higher depression scores, although the later did not survive FDR-correction (p-FDR = 0.107).

Directed metabolic connectivity in all circuits showed few differences between younger and older adults (Table S9). The *t*-tests revealed one significant age differences, with older adults having greater connectivity strength than younger adults in the inferior parietal lobule to intraparietal sulcus pathway only in the working memory circuit (p = 0.003, p-FDR = 0.022). Uncorrected trends suggested a possible reduction in other working memory pathways in older adults (e.g., ventrolateral prefrontal cortex to inferior parietal lobule and medial frontal to dorsolateral prefrontal cortex, p = 0.021 and 0.020). The angular gyrus to ventrolateral prefrontal cortex connection also tended to decrease with age (p = 0.029).

#### *4.4 Age group differences in the association between directed metabolic connectivity and cognition*

We undertook age group disaggregated correlation analyses between directed metabolic connectivity in the circuits and cognitive and affective outcomes. Age moderated the strength of some GC-cognition associations. We report these significant associations for each circuit below (see Figure S1 for significant connections; and Tables S10 for the results for all connections). Within working and episodic memory circuits, no connections survived FDR correction for younger nor older adults.

Across the circuits, several connections demonstrated significant associations with cognitive performance and affect, revealing age differences in directed metabolic information transfer. In the cognitive control network, right-hemisphere proactive control from the dorsolateral prefrontal cortex to anterior cingulate was positively associated with better switch performance in older adults (r = 0.37, p-FDR = 0.036). Although the corresponding association was not significant in younger adults, the Fisher's Z test for comparison of associations between younger and older groups was not significant (p = .810). Similarly, in the reactive circuit, the connection from the anterior insula to anterior cingulate and ventrolateral prefrontal cortex to inferior frontal junction, were linked to stop-signal RT in older adults (r = 0.45, p-FDR = 0.018; r = -0.42, p-FDR = 0.025, respectively) but not younger adults. Notably, the right ventrolateral prefrontal cortex to superior parietal lobule pathway showed a significant difference between age groups (Z = -3.02, p-FDR = 0.023), indicating that the negative association in younger adults differs markedly from the non-significant association in older adults. These results suggest that efficient information flow through proactive and reactive pathways supports behavioural performance in an age-dependent manner, with specific reactive circuits being particularly important in younger adults, albeit with a negative association with performance.

In the affective regulatory network, frontolimbic and salience circuits showed two associations between directed metabolic connectivity and depression symptoms in younger adults only. Stronger left-hemisphere anterior cingulate to caudate (r = 0.62, p-FDR < 0.001) was associated with significantly more depression symptoms in younger than older people (Z = 3.36, p-FDR = 0.008). In the right hemisphere, stronger anterior insula to ventrolateral prefrontal cortex connections were associated with more anxiety symptoms in younger adults (r = 0.43, p-FDR = 0.059). Together, these findings suggest that heightened metabolic signalling through frontolimbic loops in younger adults may reflect maladaptive information transfer underlying negative affect, whereas selective salience pathways may mediate compensatory affective regulation.

Together, these results indicate that the brain dynamically adjust directional metabolic information flow across proactive, reactive, and affective regulatory circuits, and that both efficiency and strength of these pathways are modulated by age, shaping cognitive and affective outcomes across the adult lifespan. In contrast, age effects on information transfer are not in memory circuits.

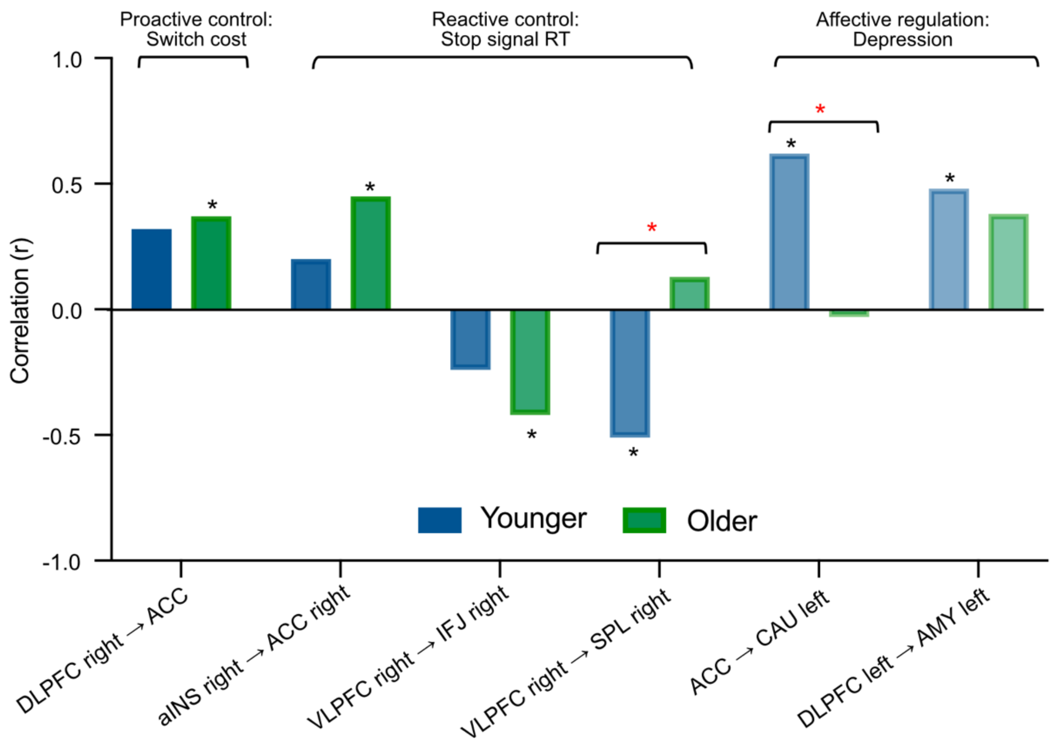


**Figure S2. Younger and older adult group Granger causality associations with cognition and affective symptomatology.** Significant correlations between directed metabolic connectivity paths in the cognitive control and memory circuits and cognitive task performance, and the affective regulatory circuit and anxiety and depression symptoms, for younger adults, older adults, or both. Black asterisk = *p-FDR < 0.05 for each age group correlation with cognition or affect; red asterisk = *p-FDR < 0.05 for difference in correlation between age groups. DLPFC = dorsolateral prefrontal cortex (BA 9/46); ACC = anterior cingulate cortex (BA 24/32); CAU = caudate; aINS = anterior insula (BA 13); VLPFC = ventrolateral prefrontal cortex (BA 44/45/47); SPL = superior parietal lobule (BA 7); AMY = amygdala. Note: all correlations are available in Table S10.

**Table S6.** Granger causality analysis for circuit connections.

| Connection | Hem | Division | N | Mean GC | SD | T-statistic | p | p-FDR | Null p | p-FDR |
|---|---|---|---|---|---|---|---|---|---|---|
| Cognitive Control | | | | | | | | | | |
| DLPFC → ACC | Left | Proactive control | 85 | 0.014 | 0.011 | 11.9 | 0.000 | 0.000 | 0.684 | 0.797 |
| DLPFC → CAU | Left | Proactive control | 84 | 0.019 | 0.018 | 9.8 | 0.000 | 0.000 | 0.013 | 0.099 |
| ACC → CAU | Left | Proactive control | 85 | 0.018 | 0.015 | 10.8 | 0.000 | 0.000 | 0.022 | 0.099 |
| ACC → DLPFC | Left | Proactive control | 84 | 0.011 | 0.012 | 8.5 | 0.000 | 0.000 | 0.990 | 0.990 |
| VLPFC → SPL | Left | Reactive control | 85 | 0.014 | 0.012 | 10.8 | 0.000 | 0.000 | 0.495 | 0.797 |
| aINS → ACC | Left | Reactive control | 84 | 0.015 | 0.013 | 10.4 | 0.000 | 0.000 | 0.212 | 0.635 |
| aINS → VLPFC | Left | Reactive control | 84 | 0.017 | 0.013 | 11.6 | 0.000 | 0.000 | 0.392 | 0.797 |
| VLPFC → IFJ | Left | Reactive control | 85 | 0.014 | 0.010 | 12.5 | 0.000 | 0.000 | 0.568 | 0.797 |
| IFJ → SPL | Left | Reactive control | 85 | 0.013 | 0.011 | 10.8 | 0.000 | 0.000 | 0.708 | 0.797 |
| DLPFC → ACC | Right | Proactive control | 85 | 0.013 | 0.011 | 10.6 | 0.000 | 0.000 | 0.684 | 0.770 |
| DLPFC → CAU | Right | Proactive control | 84 | 0.016 | 0.013 | 11.4 | 0.000 | 0.000 | 0.103 | 0.463 |
| ACC → CAU | Right | Proactive control | 85 | 0.013 | 0.013 | 9.5 | 0.000 | 0.000 | 0.644 | 0.770 |
| ACC → DLPFC | Right | Proactive control | 84 | 0.014 | 0.012 | 10.7 | 0.000 | 0.000 | 0.457 | 0.770 |
| VLPFC → SPL | Right | Reactive control | 82 | 0.015 | 0.014 | 9.5 | 0.000 | 0.000 | 0.265 | 0.770 |
| aINS → ACC | Right | Reactive control | 84 | 0.014 | 0.012 | 10.7 | 0.000 | 0.000 | 0.525 | 0.770 |
| aINS → VLPFC | Right | Reactive control | 80 | 0.012 | 0.013 | 8.1 | 0.000 | 0.000 | 0.959 | 0.959 |
| IFJ → SPL | Right | Reactive control | 84 | 0.019 | 0.016 | 10.7 | 0.000 | 0.000 | 0.004 | 0.036 |
| VLPFC → IFJ | Right | Reactive control | 82 | 0.013 | 0.014 | 8.6 | 0.000 | 0.000 | 0.617 | 0.770 |
| Working memory | | | | | | | | | | |
| IPS → DLPFC | Left | Attention-control | 84 | 0.012 | 0.010 | 10.9 | 0.000 | 0.000 | 0.901 | 0.966 |
| DLPFC → IPS | Left | Attention-control | 83 | 0.016 | 0.013 | 10.8 | 0.000 | 0.000 | 0.099 | 0.396 |
| DLPFC → VLPFC | Left | Attention-control | 84 | 0.014 | 0.012 | 10.3 | 0.000 | 0.000 | 0.632 | 0.843 |
| mPFC → DLPFC | Left | Attention-control | 84 | 0.014 | 0.013 | 10.2 | 0.000 | 0.000 | 0.364 | 0.727 |
| mPFC → IPL | Left | Attention-control | 85 | 0.018 | 0.016 | 10.2 | 0.000 | 0.000 | 0.013 | 0.104 |
| VLPFC → IPL | Left | Storage-maintenance | 84 | 0.014 | 0.012 | 10.9 | 0.000 | 0.000 | 0.486 | 0.777 |
| IPL → VLPFC | Left | Storage-maintenance | 85 | 0.015 | 0.016 | 8.6 | 0.000 | 0.000 | 0.165 | 0.440 |
| IPL → IPS | Left | Storage-maintenance | 84 | 0.012 | 0.011 | 10.4 | 0.000 | 0.000 | 0.966 | 0.966 |
| IPS → DLPFC | Right | Attention-control | 86 | 0.016 | 0.014 | 10.5 | 0.000 | 0.000 | 0.095 | 0.253 |
| DLPFC → IPS | Right | Attention-control | 86 | 0.016 | 0.013 | 11.4 | 0.000 | 0.000 | 0.072 | 0.253 |
| DLPFC → VLPFC | Right | Attention-control | 82 | 0.018 | 0.017 | 9.3 | 0.000 | 0.000 | 0.017 | 0.136 |
| mPFC → DLPFC | Right | Attention-control | 86 | 0.013 | 0.012 | 10.5 | 0.000 | 0.000 | 0.629 | 0.744 |
| mPFC → IPL | Right | Attention-control | 83 | 0.015 | 0.013 | 10.5 | 0.000 | 0.000 | 0.340 | 0.679 |
| VLPFC → IPL | Right | Storage-maintenance | 82 | 0.014 | 0.014 | 8.6 | 0.000 | 0.000 | 0.673 | 0.744 |
| IPL → VLPFC | Right | Storage-maintenance | 82 | 0.013 | 0.012 | 9.9 | 0.000 | 0.000 | 0.720 | 0.744 |
| IPL → IPS | Right | Storage-maintenance | 84 | 0.016 | 0.013 | 11.0 | 0.000 | 0.000 | 0.744 | 0.744 |
| Verbal episodic memory | | | | | | | | | | |
| VLPFC → PRC | Left | Encoding-binding | 85 | 0.014 | 0.011 | 11.6 | 0.000 | 0.000 | 0.496 | 0.592 |
| VLPFC → HIP | Left | Encoding-binding | 85 | 0.013 | 0.013 | 9.0 | 0.000 | 0.000 | 0.765 | 0.765 |
| PRC → HIP | Left | Encoding-binding | 85 | 0.014 | 0.014 | 9.1 | 0.000 | 0.000 | 0.502 | 0.592 |
| HIP → PRC | Left | Encoding-binding | 85 | 0.014 | 0.013 | 10.1 | 0.000 | 0.000 | 0.526 | 0.592 |
| PRC → pMTG | Left | Reconstruction-retrieval | 84 | 0.014 | 0.013 | 9.9 | 0.000 | 0.000 | 0.476 | 0.592 |
| HIP → pMTG | Left | Reconstruction-retrieval | 85 | 0.015 | 0.014 | 10.0 | 0.000 | 0.000 | 0.272 | 0.592 |
| pMTG → AG | Left | Reconstruction-retrieval | 85 | 0.014 | 0.012 | 10.6 | 0.000 | 0.000 | 0.455 | 0.592 |
| HIP → AG | Left | Reconstruction-retrieval | 84 | 0.015 | 0.014 | 10.0 | 0.000 | 0.000 | 0.229 | 0.592 |
| AG → VLPFC | Left | Reconstruction-retrieval | 83 | 0.016 | 0.014 | 10.2 | 0.000 | 0.000 | 0.077 | 0.592 |
| VLPFC → PRC | Right | Encoding-binding | 82 | 0.017 | 0.015 | 10.4 | 0.000 | 0.000 | 0.022 | 0.198 |
| VLPFC → HIP | Right | Encoding-binding | 81 | 0.012 | 0.011 | 9.4 | 0.000 | 0.000 | 0.909 | 0.937 |
| PRC → HIP | Right | Encoding-binding | 84 | 0.012 | 0.012 | 9.1 | 0.000 | 0.000 | 0.937 | 0.937 |
| HIP → PRC | Right | Encoding-binding | 84 | 0.014 | 0.012 | 10.5 | 0.000 | 0.000 | 0.492 | 0.885 |
| PRC → pMTG | Right | Reconstruction-retrieval | 86 | 0.016 | 0.014 | 10.3 | 0.000 | 0.000 | 0.119 | 0.535 |
| HIP → pMTG | Right | Reconstruction-retrieval | 84 | 0.014 | 0.012 | 10.9 | 0.000 | 0.000 | 0.443 | 0.885 |
| pMTG → AG | Right | Reconstruction-retrieval | 82 | 0.012 | 0.010 | 11.0 | 0.000 | 0.000 | 0.886 | 0.937 |
| HIP → AG | Right | Reconstruction-retrieval | 84 | 0.013 | 0.011 | 10.9 | 0.000 | 0.000 | 0.691 | 0.937 |
| AG → VLPFC | Right | Reconstruction-retrieval | 82 | 0.015 | 0.015 | 9.0 | 0.000 | 0.000 | 0.348 | 0.885 |
| Affective regulation | | | | | | | | | | |
| DLPFC → AMY | Left | Frontolimbic | 84 | 0.014 | 0.012 | 10.8 | 0.000 | 0.000 | 0.421 | 0.601 |
| VLPFC → AMY | Left | Frontolimbic | 85 | 0.012 | 0.011 | 10.8 | 0.000 | 0.000 | 0.889 | 0.889 |
| AMY → DLPFC | Left | Frontolimbic | 84 | 0.015 | 0.013 | 11.1 | 0.000 | 0.000 | 0.180 | 0.424 |
| ACC → CAU | Left | Frontolimbic | 85 | 0.018 | 0.015 | 10.8 | 0.000 | 0.000 | 0.022 | 0.110 |
| ACC → AMY | Left | Frontolimbic | 85 | 0.013 | 0.011 | 10.3 | 0.000 | 0.000 | 0.759 | 0.844 |
| DLPFC → CAU | Left | Frontolimbic | 84 | 0.019 | 0.018 | 9.8 | 0.000 | 0.000 | 0.013 | 0.110 |
| AMY → aINS | Left | Salience | 83 | 0.013 | 0.010 | 11.5 | 0.000 | 0.000 | 0.721 | 0.844 |
| ACC → aINS | Left | Salience | 85 | 0.017 | 0.013 | 12.4 | 0.000 | 0.000 | 0.042 | 0.140 |
| aINS → VLPFC | Left | Salience | 84 | 0.017 | 0.013 | 11.6 | 0.000 | 0.000 | 0.392 | 0.601 |
| aINS → ACC | Left | Salience | 84 | 0.015 | 0.013 | 10.4 | 0.000 | 0.000 | 0.212 | 0.424 |
| AMY → DLPFC | Right | Frontolimbic | 85 | 0.015 | 0.012 | 11.8 | 0.000 | 0.000 | 0.192 | 0.639 |
| DLPFC → CAU | Right | Frontolimbic | 84 | 0.016 | 0.013 | 11.4 | 0.000 | 0.000 | 0.103 | 0.584 |
| VLPFC → AMY | Right | Frontolimbic | 82 | 0.014 | 0.012 | 10.4 | 0.000 | 0.000 | 0.466 | 0.751 |
| ACC → AMY | Right | Frontolimbic | 84 | 0.014 | 0.012 | 10.3 | 0.000 | 0.000 | 0.450 | 0.751 |
| ACC → CAU | Right | Frontolimbic | 85 | 0.013 | 0.013 | 9.5 | 0.000 | 0.000 | 0.644 | 0.805 |
| DLPFC → AMY | Right | Frontolimbic | 85 | 0.013 | 0.014 | 8.5 | 0.000 | 0.000 | 0.730 | 0.811 |
| aINS → ACC | Right | Salience | 84 | 0.014 | 0.012 | 10.7 | 0.000 | 0.000 | 0.525 | 0.751 |
| AMY → aINS | Right | Salience | 85 | 0.016 | 0.015 | 9.9 | 0.000 | 0.000 | 0.117 | 0.584 |
| ACC → aINS | Right | Salience | 85 | 0.014 | 0.014 | 9.5 | 0.000 | 0.000 | 0.496 | 0.751 |
| aINS → VLPFC | Right | Salience | 80 | 0.012 | 0.013 | 8.1 | 0.000 | 0.000 | 0.959 | 0.959 |

**Table S7.** Correlation of Granger causality with cognition and behaviour measures.

| Connection | Division | Hem | Measure | N | Correlation (r) | p-value | p-FDR |
|---|---|---|---|---|---|---|---|
| Cognitive control | | | | | | | |
| DLPFC → ACC | Proactive | Left | Switch cost (reversed) | 85 | 0.19 | 0.075 | 0.300 |
| DLPFC → CAU | Proactive | Left | Switch cost (reversed) | 85 | -0.10 | 0.380 | 0.507 |
| ACC → CAU | Proactive | Left | Switch cost (reversed) | 85 | 0.13 | 0.228 | 0.456 |
| ACC → DLPFC | Proactive | Left | Switch cost (reversed) | 83 | -0.04 | 0.736 | 0.736 |
| DLPFC → ACC | Proactive | Right | Switch cost (reversed) | 85 | 0.32 | 0.002 | 0.010 |
| DLPFC → CAU | Proactive | Right | Switch cost (reversed) | 84 | 0.22 | 0.044 | 0.087 |
| ACC → CAU | Proactive | Right | Switch cost (reversed) | 84 | 0.02 | 0.874 | 0.874 |
| ACC → DLPFC | Proactive | Right | Switch cost (reversed) | 83 | -0.16 | 0.146 | 0.195 |
| VLPFC → SPL | Reactive | Left | Stop signal RT (reversed) | 85 | -0.01 | 0.897 | 0.951 |
| aINS → ACC | Reactive | Left | Stop signal RT (reversed) | 85 | 0.05 | 0.647 | 0.951 |
| aINS → VLPFC | Reactive | Left | Stop signal RT (reversed) | 85 | -0.01 | 0.951 | 0.951 |
| VLPFC → IFJ | Reactive | Left | Stop signal RT (reversed) | 85 | 0.05 | 0.647 | 0.951 |
| IFJ → SPL | Reactive | Left | Stop signal RT (reversed) | 85 | -0.11 | 0.326 | 0.951 |
| VLPFC → SPL | Reactive | Right | Stop signal RT (reversed) | 82 | -0.21 | 0.053 | 0.088 |
| aINS → ACC | Reactive | Right | Stop signal RT (reversed) | 85 | 0.30 | 0.005 | 0.012 |
| aINS → VLPFC | Reactive | Right | Stop signal RT (reversed) | 81 | -0.05 | 0.645 | 0.772 |
| VLPFC → IFJ | Reactive | Right | Stop signal RT (reversed) | 82 | -0.31 | 0.005 | 0.012 |
| IFJ → SPL | Reactive | Right | Stop signal RT (reversed) | 84 | 0.03 | 0.772 | 0.772 |
| Working memory | | | | | | | |
| VLPFC → IPL | Storage-maintenance | Left | Digit span total | 84 | 0.00 | 0.976 | 0.976 |
| IPL → VLPFC | Storage-maintenance | Left | Digit span total | 85 | 0.15 | 0.180 | 0.719 |
| IPL → IPS | Storage-maintenance | Left | Digit span total | 84 | 0.22 | 0.047 | 0.376 |
| IPS → DLPFC | Attention-control | Left | Digit span total | 84 | -0.09 | 0.425 | 0.849 |
| DLPFC → IPS | Attention-control | Left | Digit span total | 83 | -0.07 | 0.531 | 0.849 |
| DLPFC → VLPFC | Attention-control | Left | Digit span total | 85 | 0.03 | 0.806 | 0.976 |
| mPFC → DLPFC | Attention-control | Left | Digit span total | 84 | -0.10 | 0.385 | 0.849 |
| mPFC → IPL | Attention-control | Left | Digit span total | 85 | 0.02 | 0.863 | 0.976 |
| VLPFC → IPL | Storage-maintenance | Right | Digit span total | 81 | -0.03 | 0.822 | 0.822 |
| IPL → VLPFC | Storage-maintenance | Right | Digit span total | 82 | 0.07 | 0.514 | 0.822 |
| IPL → IPS | Storage-maintenance | Right | Digit span total | 86 | 0.06 | 0.598 | 0.822 |
| IPS → DLPFC | Attention-control | Right | Digit span total | 86 | 0.03 | 0.792 | 0.822 |
| DLPFC → IPS | Attention-control | Right | Digit span total | 86 | -0.29 | 0.007 | 0.055 |
| DLPFC → VLPFC | Attention-control | Right | Digit span total | 82 | 0.05 | 0.668 | 0.822 |
| mPFC → DLPFC | Attention-control | Right | Digit span total | 84 | -0.16 | 0.152 | 0.607 |
| mPFC → IPL | Attention-control | Right | Digit span total | 84 | 0.06 | 0.566 | 0.822 |
| Verbal episodic memory | | | | | | | |
| VLPFC → PRC | Encoding-binding | Left | HVLT delayed recall | 86 | -0.08 | 0.445 | 0.883 |
| VLPFC → HIP | Encoding-binding | Left | HVLT delayed recall | 84 | 0.06 | 0.605 | 0.883 |
| PRC → HIP | Encoding-binding | Left | HVLT delayed recall | 85 | 0.14 | 0.216 | 0.883 |
| HIP → PRC | Encoding-binding | Left | HVLT delayed recall | 85 | 0.29 | 0.007 | 0.062 |
| PRC → pMTG | Reconstruction-rertieval | Left | HVLT delayed recall | 84 | 0.02 | 0.883 | 0.883 |
| HIP → pMTG | Reconstruction-rertieval | Left | HVLT delayed recall | 85 | 0.09 | 0.393 | 0.883 |
| pMTG → AG | Reconstruction-rertieval | Left | HVLT delayed recall | 85 | -0.03 | 0.791 | 0.883 |
| HIP → AG | Reconstruction-rertieval | Left | HVLT delayed recall | 84 | -0.02 | 0.827 | 0.883 |
| AG → VLPFC | Reconstruction-rertieval | Left | HVLT delayed recall | 83 | 0.04 | 0.705 | 0.883 |
| VLPFC → PRC | Encoding-binding | Right | HVLT delayed recall | 82 | -0.09 | 0.404 | 0.921 |
| VLPFC → HIP | Encoding-binding | Right | HVLT delayed recall | 81 | -0.01 | 0.921 | 0.921 |
| PRC → HIP | Encoding-binding | Right | HVLT delayed recall | 84 | -0.08 | 0.490 | 0.921 |
| HIP → PRC | Encoding-binding | Right | HVLT delayed recall | 84 | -0.04 | 0.688 | 0.921 |
| PRC → pMTG | Reconstruction-rertieval | Right | HVLT delayed recall | 86 | -0.02 | 0.880 | 0.921 |
| HIP → pMTG | Reconstruction-rertieval | Right | HVLT delayed recall | 84 | 0.03 | 0.760 | 0.921 |
| pMTG → AG | Reconstruction-rertieval | Right | HVLT delayed recall | 84 | -0.09 | 0.422 | 0.921 |
| HIP → AG | Reconstruction-rertieval | Right | HVLT delayed recall | 84 | -0.14 | 0.194 | 0.921 |
| AG → VLPFC | Reconstruction-rertieval | Right | HVLT delayed recall | 81 | -0.03 | 0.779 | 0.921 |

*Continued …*

*Table S7 continued …*

| Connection | Division | Hem | Measure | N | Correlation (r) | p-value | p-FDR |
|---|---|---|---|---|---|---|---|
| Affective regulatory | | | | | | | |
| DLPFC → AMY | Frontolimbic | Left | Beck anxiety | 83 | 0.34 | 0.002 | 0.018 |
| VLPFC → AMY | Frontolimbic | Left | Beck anxiety | 84 | 0.18 | 0.104 | 0.271 |
| ACC → AMY | Frontolimbic | Left | Beck anxiety | 83 | -0.05 | 0.655 | 0.971 |
| AMY → DLPFC | Frontolimbic | Left | Beck anxiety | 83 | 0.18 | 0.108 | 0.271 |
| ACC → CAU | Frontolimbic | Left | Beck anxiety | 84 | 0.00 | 0.971 | 0.971 |
| DLPFC → CAU | Frontolimbic | Left | Beck anxiety | 84 | -0.05 | 0.637 | 0.971 |
| AMY → aINS | Salience | Left | Beck anxiety | 83 | 0.04 | 0.699 | 0.971 |
| aINS → ACC | Salience | Left | Beck anxiety | 84 | -0.02 | 0.866 | 0.971 |
| ACC → aINS | Salience | Left | Beck anxiety | 83 | -0.18 | 0.097 | 0.271 |
| aINS → VLPFC | Salience | Left | Beck anxiety | 84 | 0.02 | 0.889 | 0.971 |
| DLPFC → AMY | Frontolimbic | Right | Beck anxiety | 83 | 0.19 | 0.085 | 0.244 |
| VLPFC → AMY | Frontolimbic | Right | Beck anxiety | 80 | 0.06 | 0.625 | 0.695 |
| ACC → AMY | Frontolimbic | Right | Beck anxiety | 84 | -0.06 | 0.605 | 0.695 |
| AMY → DLPFC | Frontolimbic | Right | Beck anxiety | 84 | -0.12 | 0.262 | 0.523 |
| ACC → CAU | Frontolimbic | Right | Beck anxiety | 83 | 0.07 | 0.546 | 0.695 |
| DLPFC → CAU | Frontolimbic | Right | Beck anxiety | 83 | -0.18 | 0.097 | 0.244 |
| AMY → aINS | Salience | Right | Beck anxiety | 85 | 0.18 | 0.097 | 0.244 |
| aINS → ACC | Salience | Right | Beck anxiety | 84 | 0.02 | 0.881 | 0.881 |
| ACC → aINS | Salience | Right | Beck anxiety | 84 | -0.06 | 0.598 | 0.695 |
| aINS → VLPFC | Salience | Right | Beck anxiety | 80 | 0.41 | 0.000 | 0.002 |
| DLPFC → AMY | Frontolimbic | Left | Beck depression | 83 | 0.46 | 0.000 | 0.000 |
| VLPFC → AMY | Frontolimbic | Left | Beck depression | 84 | 0.01 | 0.935 | 0.935 |
| ACC → AMY | Frontolimbic | Left | Beck depression | 83 | -0.11 | 0.325 | 0.752 |
| AMY → DLPFC | Frontolimbic | Left | Beck depression | 83 | 0.06 | 0.618 | 0.843 |
| ACC → CAU | Frontolimbic | Left | Beck depression | 84 | 0.26 | 0.015 | 0.076 |
| DLPFC → CAU | Frontolimbic | Left | Beck depression | 84 | -0.04 | 0.697 | 0.843 |
| AMY → aINS | Salience | Left | Beck depression | 83 | 0.03 | 0.758 | 0.843 |
| aINS → ACC | Salience | Left | Beck depression | 84 | 0.10 | 0.376 | 0.752 |
| ACC → aINS | Salience | Left | Beck depression | 83 | -0.24 | 0.027 | 0.091 |
| aINS → VLPFC | Salience | Left | Beck depression | 84 | -0.06 | 0.565 | 0.843 |
| DLPFC → AMY | Frontolimbic | Right | Beck depression | 83 | 0.08 | 0.474 | 0.641 |
| VLPFC → AMY | Frontolimbic | Right | Beck depression | 80 | -0.04 | 0.744 | 0.744 |
| ACC → AMY | Frontolimbic | Right | Beck depression | 84 | -0.10 | 0.348 | 0.641 |
| AMY → DLPFC | Frontolimbic | Right | Beck depression | 84 | -0.04 | 0.708 | 0.744 |
| ACC → CAU | Frontolimbic | Right | Beck depression | 83 | -0.08 | 0.495 | 0.641 |
| DLPFC → CAU | Frontolimbic | Right | Beck depression | 83 | -0.15 | 0.186 | 0.641 |
| AMY → aINS | Salience | Right | Beck depression | 85 | 0.16 | 0.155 | 0.641 |
| aINS → ACC | Salience | Right | Beck depression | 84 | 0.07 | 0.511 | 0.641 |
| ACC → aINS | Salience | Right | Beck depression | 84 | 0.07 | 0.513 | 0.641 |
| aINS → VLPFC | Salience | Right | Beck depression | 80 | 0.08 | 0.470 | 0.641 |

**Table S8.** Mean and standard deviation of cognitive and behaviour measures for younger and older adult groups, and *t*-test of age group differences (p, two-sided).

| | Younger adults | | Older adults | | Younger vs older | | |
|---|---|---|---|---|---|---|---|
| | Mean | SD | Mean | SD | t-value | p | p-FDR |
| HVLT: Delayed recall | 9.3 | 2.5 | 7.3 | 2.6 | 3.7 | 0.000 | 0.000 |
| Digit span: Total longest forward and backward | 12.3 | 2.7 | 11.9 | 1.8 | 0.8 | 0.449 | 0.449 |
| Category switch: Switch cost (sec) | 0.28 | 0.23 | 2.13 | 0.48 | 0.4 | 0.011 | 0.017 |
| Stop signal: RT in SS trials (sec) | 0.53 | 0.13 | 0.60 | 0.12 | 2.6 | 0.010 | 0.017 |
| Beck depression inventory | 8.6 | 8.4 | 4.2 | 4.5 | 3.0 | 0.003 | 0.009 |
| Beck anxiety inventory | 6.8 | 8.8 | 4.3 | 3.8 | 1.7 | 0.088 | 0.106 |

**Table S9.** Mean and standard deviation of Granger causality values for younger and older adult groups, and t-test of age group differences (p two-sided).

| Connection | Division | Hem | Younger N | Younger Mean | Older N | Older Mean | t | df | p | p-FDR |
|---|---|---|---|---|---|---|---|---|---|---|
| Cogntive control | | | | | | | | | | |
| ACC → Caudate | Proactive | Left | 40 | 0.017 | 46 | 0.017 | 0.112 | 76 | 0.911 | 0.955 |
| ACC → DLPFC | Proactive | Left | 39 | 0.010 | 45 | 0.014 | -1.680 | 72 | 0.097 | 0.409 |
| DLPFC → ACC | Proactive | Left | 39 | 0.018 | 46 | 0.011 | 1.507 | 69 | 0.136 | 0.409 |
| DLPFC → Caudate | Proactive | Left | 39 | 0.020 | 45 | 0.021 | -0.290 | 81 | 0.773 | 0.955 |
| IFJ → PPC | Reactive | Left | 40 | 0.011 | 45 | 0.014 | -0.958 | 81 | 0.341 | 0.614 |
| VLPFC → IFJ | Reactive | Left | 40 | 0.013 | 45 | 0.016 | -1.179 | 83 | 0.242 | 0.544 |
| VLPFC → PPC | Reactive | Left | 39 | 0.016 | 46 | 0.014 | -0.056 | 82 | 0.955 | 0.955 |
| aINS → ACC | Reactive | Left | 40 | 0.015 | 45 | 0.015 | 0.733 | 82 | 0.466 | 0.699 |
| aINS → VLPFC | Reactive | Left | 39 | 0.015 | 46 | 0.020 | -1.899 | 83 | 0.061 | 0.409 |
| ACC → Caudate | Proactive | Right | 39 | 0.011 | 45 | 0.016 | -1.794 | 63 | 0.078 | 0.370 |
| ACC → DLPFC | Proactive | Right | 40 | 0.012 | 45 | 0.018 | -1.762 | 74 | 0.082 | 0.370 |
| DLPFC → ACC | Proactive | Right | 40 | 0.013 | 45 | 0.015 | -0.438 | 82 | 0.663 | 0.874 |
| DLPFC → Caudate | Proactive | Right | 39 | 0.015 | 45 | 0.019 | -1.102 | 81 | 0.274 | 0.804 |
| IFJ → PPC | Reactive | Right | 38 | 0.023 | 46 | 0.018 | 0.284 | 80 | 0.777 | 0.874 |
| VLPFC → IFJ | Reactive | Right | 40 | 0.013 | 42 | 0.015 | -0.059 | 77 | 0.953 | 0.953 |
| VLPFC → PPC | Reactive | Right | 40 | 0.014 | 42 | 0.018 | -0.516 | 79 | 0.607 | 0.874 |
| aINS → ACC | Reactive | Right | 39 | 0.017 | 45 | 0.014 | 0.926 | 78 | 0.358 | 0.804 |
| aINS → VLPFC | Reactive | Right | 39 | 0.016 | 43 | 0.013 | 0.377 | 75 | 0.707 | 0.874 |
| Working memory | | | | | | | | | | |
| DLPFC → IPS | Attention-control | Left | 38 | 0.020 | 45 | 0.018 | 0.103 | 81 | 0.918 | 0.918 |
| DLPFC → VLPFC | Attention-control | Left | 40 | 0.015 | 45 | 0.014 | 0.650 | 72 | 0.517 | 0.591 |
| IPS → DLPFC | Attention-control | Left | 39 | 0.011 | 45 | 0.013 | -0.927 | 81 | 0.357 | 0.549 |
| mPFC → DLPFC | Attention-control | Left | 39 | 0.013 | 45 | 0.017 | -1.628 | 81 | 0.107 | 0.215 |
| mPFC → IPL | Attention-control | Left | 39 | 0.016 | 46 | 0.020 | -1.785 | 82 | 0.078 | 0.208 |
| IPL → IPS | Storage-maintenance | Left | 39 | 0.009 | 45 | 0.017 | -3.104 | 66 | 0.003 | 0.022 |
| IPL → VLPFC | Storage-maintenance | Left | 40 | 0.017 | 45 | 0.015 | 0.826 | 70 | 0.412 | 0.549 |
| VLPFC → IPL | Storage-maintenance | Left | 39 | 0.018 | 45 | 0.012 | 2.362 | 69 | 0.021 | 0.084 |
| DLPFC → IPS | Attention-control | Right | 39 | 0.015 | 46 | 0.016 | -1.054 | 83 | 0.295 | 0.505 |
| DLPFC → VLPFC | Attention-control | Right | 40 | 0.015 | 43 | 0.019 | -1.049 | 77 | 0.297 | 0.505 |
| IPS → DLPFC | Attention-control | Right | 40 | 0.017 | 46 | 0.014 | 0.811 | 76 | 0.420 | 0.560 |
| mPFC → DLPFC | Attention-control | Right | 40 | 0.016 | 44 | 0.012 | 2.367 | 76 | 0.020 | 0.164 |
| mPFC → IPL | Attention-control | Right | 39 | 0.015 | 45 | 0.015 | -0.168 | 80 | 0.867 | 0.867 |
| IPL → IPS | Storage-maintenance | Right | 40 | 0.017 | 46 | 0.016 | 0.206 | 82 | 0.837 | 0.867 |
| IPL → VLPFC | Storage-maintenance | Right | 39 | 0.016 | 43 | 0.016 | -1.269 | 80 | 0.208 | 0.505 |
| VLPFC → IPL | Storage-maintenance | Right | 38 | 0.018 | 42 | 0.012 | 1.010 | 74 | 0.316 | 0.505 |
| Verbal episodic memory | | | | | | | | | | |
| HIP → PRC | Encoding-binding | Left | 39 | 0.017 | 46 | 0.013 | 0.874 | 83 | 0.384 | 0.849 |
| PRC → HIP | Encoding-binding | Left | 39 | 0.014 | 46 | 0.015 | -0.949 | 80 | 0.345 | 0.849 |
| VLPFC → HIP | Encoding-binding | Left | 39 | 0.013 | 45 | 0.014 | -0.314 | 81 | 0.755 | 0.849 |
| VLPFC → PRC | Encoding-binding | Left | 39 | 0.014 | 46 | 0.016 | -1.193 | 82 | 0.236 | 0.849 |
| AG → VLPFC | Reconstruction-retrieval | Left | 39 | 0.020 | 44 | 0.018 | 0.792 | 64 | 0.432 | 0.849 |
| HIP → AG | Reconstruction-retrieval | Left | 39 | 0.017 | 45 | 0.017 | 0.007 | 82 | 0.994 | 0.994 |
| HIP → pMTG | Reconstruction-retrieval | Left | 39 | 0.016 | 46 | 0.016 | -0.427 | 79 | 0.671 | 0.849 |
| PRC → pMTG | Reconstruction-retrieval | Left | 39 | 0.015 | 45 | 0.016 | -0.364 | 82 | 0.717 | 0.849 |
| pMTG → AG | Reconstruction-retrieval | Left | 40 | 0.014 | 45 | 0.016 | -0.404 | 83 | 0.687 | 0.849 |
| HIP → PRC | Encoding-binding | Right | 40 | 0.012 | 45 | 0.018 | -1.658 | 68 | 0.102 | 0.458 |
| PRC → HIP | Encoding-binding | Right | 39 | 0.012 | 45 | 0.014 | -0.764 | 80 | 0.447 | 0.701 |
| VLPFC → HIP | Encoding-binding | Right | 39 | 0.012 | 41 | 0.016 | -0.585 | 74 | 0.560 | 0.720 |
| VLPFC → PRC | Encoding-binding | Right | 40 | 0.020 | 43 | 0.017 | 0.939 | 75 | 0.351 | 0.701 |
| AG → VLPFC | Reconstruction-retrieval | Right | 39 | 0.018 | 41 | 0.012 | 2.231 | 61 | 0.029 | 0.264 |
| HIP → AG | Reconstruction-retrieval | Right | 40 | 0.015 | 44 | 0.014 | 1.013 | 76 | 0.314 | 0.701 |
| HIP → pMTG | Reconstruction-retrieval | Right | 39 | 0.016 | 45 | 0.015 | 0.046 | 80 | 0.963 | 0.963 |
| PRC → pMTG | Reconstruction-retrieval | Right | 40 | 0.014 | 46 | 0.016 | -0.730 | 84 | 0.468 | 0.701 |
| pMTG → AG | Reconstruction-retrieval | Right | 39 | 0.014 | 45 | 0.015 | -0.144 | 76 | 0.886 | 0.963 |
| Cogntive control | | | | | | | | | | |
| ACC → Amygdala | Frontolimbic | Left | 40 | 0.013 | 45 | 0.014 | -0.020 | 83 | 0.984 | 0.984 |
| ACC → Caudate | Frontolimbic | Left | 40 | 0.017 | 46 | 0.017 | 0.112 | 76 | 0.911 | 0.984 |
| Amygdala → DLPFC | Frontolimbic | Left | 39 | 0.018 | 45 | 0.016 | 0.845 | 81 | 0.401 | 0.931 |
| DLPFC → Amygdala | Frontolimbic | Left | 40 | 0.016 | 44 | 0.015 | 1.435 | 73 | 0.156 | 0.778 |
| DLPFC → Caudate | Frontolimbic | Left | 39 | 0.020 | 45 | 0.021 | -0.290 | 81 | 0.773 | 0.984 |
| VLPFC → Amygdala | Frontolimbic | Left | 39 | 0.012 | 45 | 0.014 | -0.567 | 81 | 0.573 | 0.954 |
| ACC → aINS | Salience | Left | 39 | 0.016 | 46 | 0.018 | -1.187 | 82 | 0.239 | 0.796 |
| Amygdala → aINS | Salience | Left | 39 | 0.014 | 45 | 0.015 | -0.269 | 81 | 0.788 | 0.984 |
| aINS → ACC | Salience | Left | 40 | 0.015 | 45 | 0.015 | 0.733 | 82 | 0.466 | 0.931 |
| aINS → VLPFC | Salience | Left | 39 | 0.015 | 46 | 0.020 | -1.899 | 83 | 0.061 | 0.610 |
| ACC → Amygdala | Frontolimbic | Right | 39 | 0.013 | 45 | 0.016 | -1.042 | 79 | 0.301 | 0.596 |
| ACC → Caudate | Frontolimbic | Right | 39 | 0.011 | 45 | 0.016 | -1.794 | 63 | 0.078 | 0.596 |
| Amygdala → DLPFC | Frontolimbic | Right | 40 | 0.015 | 46 | 0.016 | -0.217 | 83 | 0.829 | 0.890 |
| DLPFC → Amygdala | Frontolimbic | Right | 39 | 0.014 | 45 | 0.014 | 0.139 | 68 | 0.890 | 0.890 |
| DLPFC → Caudate | Frontolimbic | Right | 39 | 0.015 | 45 | 0.019 | -1.102 | 81 | 0.274 | 0.596 |
| VLPFC → Amygdala | Frontolimbic | Right | 39 | 0.013 | 42 | 0.015 | -0.591 | 78 | 0.556 | 0.795 |
| ACC → aINS | Salience | Right | 40 | 0.017 | 46 | 0.013 | 1.030 | 70 | 0.306 | 0.596 |
| Amygdala → aINS | Salience | Right | 40 | 0.019 | 45 | 0.015 | 1.377 | 65 | 0.173 | 0.596 |
| aINS → ACC | Salience | Right | 39 | 0.017 | 45 | 0.014 | 0.926 | 78 | 0.358 | 0.596 |
| aINS → VLPFC | Salience | Right | 39 | 0.016 | 43 | 0.013 | 0.377 | 75 | 0.707 | 0.884 |

**Table S10.** Correlation of Granger causality in cognitive control circuit with cognitive test performance for younger and older adult groups.

| Directed connectivity | | | | Younger adults | | | | Older adults | | | | Younger vs older adults | | |
|---|---|---|---|---|---|---|---|---|---|---|---|---|---|---|
| Connection | Division | Hem | Measure | r | p | p-FDR | N | r | p | p-FDR | N | Z | p | p-FDR |
| Cognitive control | | | | | | | | | | | | | | |
| ACC → CAU_left | Proactive | Left | Switch cost | 0.02 | 0.892 | 0.892 | 40 | 0.21 | 0.160 | 0.898 | 46 | -0.85 | 0.393 | 0.927 |
| ACC → DLPFC_left | Proactive | Left | Switch cost | 0.08 | 0.636 | 0.768 | 39 | 0.02 | 0.878 | 0.946 | 45 | 0.24 | 0.809 | 0.927 |
| DLPFC → ACC_left | Proactive | Left | Switch cost | 0.33 | 0.039 | 0.354 | 39 | 0.07 | 0.626 | 0.898 | 46 | 1.20 | 0.231 | 0.927 |
| DLPFC → CAU_left | Proactive | Left | Switch cost | -0.12 | 0.481 | 0.768 | 39 | -0.07 | 0.666 | 0.898 | 45 | -0.22 | 0.824 | 0.927 |
| aINS → ACC_left | Reactive | Left | Stop signal RT | 0.07 | 0.651 | 0.768 | 40 | -0.01 | 0.946 | 0.946 | 45 | 0.37 | 0.709 | 0.927 |
| aINS → VLPFC_left | Reactive | Left | Stop signal RT | -0.07 | 0.674 | 0.768 | 39 | 0.15 | 0.327 | 0.898 | 46 | -0.97 | 0.333 | 0.927 |
| IFJ → SPL_left | Reactive | Left | Stop signal RT | -0.09 | 0.586 | 0.768 | 40 | -0.08 | 0.621 | 0.898 | 45 | -0.06 | 0.954 | 0.954 |
| VLPFC → IFJ_left | Reactive | Left | Stop signal RT | 0.12 | 0.445 | 0.768 | 40 | 0.06 | 0.699 | 0.898 | 45 | 0.29 | 0.771 | 0.927 |
| VLPFC → SPL_left | Reactive | Left | Stop signal RT | 0.07 | 0.683 | 0.768 | 39 | -0.08 | 0.584 | 0.898 | 46 | 0.67 | 0.504 | 0.927 |
| ACC → CAU_right | Proactive | Right | Switch cost | 0.03 | 0.841 | 0.841 | 39 | 0.07 | 0.635 | 0.715 | 45 | -0.17 | 0.861 | 0.861 |
| ACC → DLPFC_right | Proactive | Right | Switch cost | -0.31 | 0.054 | 0.162 | 40 | 0.05 | 0.756 | 0.756 | 45 | -1.62 | 0.106 | 0.317 |
| DLPFC → ACC_right | Proactive | Right | Switch cost | 0.32 | 0.042 | 0.162 | 40 | 0.37 | 0.012 | 0.036 | 45 | -0.24 | 0.810 | 0.861 |
| DLPFC → CAU_right | Proactive | Right | Switch cost | 0.20 | 0.234 | 0.301 | 39 | 0.11 | 0.482 | 0.620 | 45 | 0.40 | 0.693 | 0.861 |
| aINS → ACC_right | Reactive | Right | Stop signal RT | 0.20 | 0.217 | 0.301 | 39 | 0.45 | 0.002 | 0.018 | 45 | -1.22 | 0.222 | 0.405 |
| aINS → VLPFC_right | Reactive | Right | Stop signal RT | 0.12 | 0.474 | 0.533 | 39 | -0.16 | 0.310 | 0.558 | 43 | 1.21 | 0.225 | 0.405 |
| IFJ → SPL_right | Reactive | Right | Stop signal RT | -0.23 | 0.173 | 0.301 | 38 | 0.24 | 0.116 | 0.260 | 46 | -2.06 | 0.039 | 0.176 |
| VLPFC → IFJ_right | Reactive | Right | Stop signal RT | -0.24 | 0.139 | 0.301 | 40 | -0.42 | 0.006 | 0.025 | 42 | 0.89 | 0.372 | 0.558 |
| VLPFC → SPL_right | Reactive | Right | Stop signal RT | -0.51 | 0.001 | 0.007 | 40 | 0.13 | 0.417 | 0.620 | 42 | -3.02 | 0.003 | 0.023 |
| Working memory | | | | | | | | | | | | | | |
| DLPFC → IPS_left | Attention-control | Left | Digit span total | -0.19 | 0.265 | 0.424 | 38 | 0.05 | 0.732 | 0.969 | 45 | -1.05 | 0.294 | 0.470 |
| DLPFC → VLPFC_left | Attention-control | Left | Digit span total | 0.11 | 0.491 | 0.561 | 40 | -0.15 | 0.315 | 0.969 | 45 | 1.18 | 0.236 | 0.470 |
| IPS → DLPFC_left | Attention-control | Left | Digit span total | -0.27 | 0.095 | 0.189 | 39 | 0.07 | 0.632 | 0.969 | 45 | -1.55 | 0.121 | 0.363 |
| mPFC → DLPFC_left | Attention-control | Left | Digit span total | -0.28 | 0.082 | 0.189 | 39 | 0.11 | 0.462 | 0.969 | 45 | -1.77 | 0.076 | 0.363 |
| mPFC → IPL_left | Attention-control | Left | Digit span total | 0.12 | 0.472 | 0.561 | 39 | -0.05 | 0.751 | 0.969 | 46 | 0.74 | 0.459 | 0.612 |
| IPL → IPS_left | Storage-maintenance | Left | Digit span total | 0.27 | 0.092 | 0.189 | 39 | 0.18 | 0.234 | 0.969 | 45 | 0.43 | 0.668 | 0.763 |
| IPL → VLPFC_left | Storage-maintenance | Left | Digit span total | 0.32 | 0.045 | 0.189 | 40 | -0.01 | 0.969 | 0.969 | 45 | 1.49 | 0.136 | 0.363 |
| VLPFC → IPL_left | Storage-maintenance | Left | Digit span total | -0.04 | 0.810 | 0.810 | 39 | -0.01 | 0.952 | 0.969 | 45 | -0.13 | 0.893 | 0.893 |
| DLPFC → IPS_right | Attention-control | Right | Digit span total | -0.39 | 0.015 | 0.119 | 39 | -0.09 | 0.550 | 0.883 | 46 | -1.41 | 0.159 | 0.637 |
| DLPFC → VLPFC_right | Attention-control | Right | Digit span total | 0.14 | 0.397 | 0.918 | 40 | 0.01 | 0.969 | 0.969 | 43 | 0.58 | 0.561 | 0.864 |
| IPS → DLPFC_right | Attention-control | Right | Digit span total | 0.00 | 0.987 | 0.987 | 40 | 0.06 | 0.689 | 0.883 | 46 | -0.28 | 0.777 | 0.864 |
| mPFC → DLPFC_right | Attention-control | Right | Digit span total | -0.25 | 0.120 | 0.479 | 40 | -0.09 | 0.569 | 0.883 | 44 | -0.74 | 0.462 | 0.864 |
| mPFC → IPL_right | Attention-control | Right | Digit span total | -0.07 | 0.679 | 0.918 | 39 | 0.25 | 0.100 | 0.797 | 45 | -1.42 | 0.156 | 0.637 |
| IPL → IPS_right | Storage-maintenance | Right | Digit span total | 0.04 | 0.803 | 0.918 | 40 | 0.08 | 0.602 | 0.883 | 46 | -0.17 | 0.864 | 0.864 |
| IPL → VLPFC_right | Storage-maintenance | Right | Digit span total | 0.12 | 0.461 | 0.918 | 39 | 0.05 | 0.772 | 0.883 | 43 | 0.33 | 0.738 | 0.864 |
| VLPFC → IPL_right | Storage-maintenance | Right | Digit span total | 0.06 | 0.724 | 0.918 | 38 | -0.18 | 0.267 | 0.883 | 42 | 1.01 | 0.310 | 0.828 |
| Verbal episodic memory | | | | | | | | | | | | | | |
| HIP → PRC_left | Encoding-binding | Left | HVLT delayed recall | 0.13 | 0.420 | 0.792 | 39 | 0.38 | 0.009 | 0.081 | 46 | -1.18 | 0.236 | 0.484 |
| PRC → HIP_left | Encoding-binding | Left | HVLT delayed recall | -0.04 | 0.792 | 0.792 | 39 | 0.32 | 0.029 | 0.132 | 46 | -1.67 | 0.095 | 0.286 |
| VLPFC → HIP_left | Encoding-binding | Left | HVLT delayed recall | 0.05 | 0.741 | 0.792 | 39 | 0.26 | 0.088 | 0.259 | 45 | -0.92 | 0.359 | 0.539 |
| VLPFC → PRC_left | Encoding-binding | Left | HVLT delayed recall | 0.07 | 0.688 | 0.792 | 39 | -0.18 | 0.228 | 0.342 | 46 | 1.11 | 0.269 | 0.484 |
| AG → VLPFC_left | Reconstruction-retrieval | Left | HVLT delayed recall | -0.18 | 0.273 | 0.792 | 39 | 0.20 | 0.200 | 0.342 | 44 | -1.67 | 0.095 | 0.286 |
| HIP → AG_left | Reconstruction-retrieval | Left | HVLT delayed recall | -0.09 | 0.588 | 0.792 | 39 | 0.02 | 0.883 | 0.883 | 45 | -0.49 | 0.621 | 0.621 |
| HIP → pMTG_left | Reconstruction-retrieval | Left | HVLT delayed recall | 0.21 | 0.210 | 0.792 | 39 | 0.05 | 0.746 | 0.883 | 46 | 0.70 | 0.481 | 0.618 |
| pMTG → AG_left | Reconstruction-retrieval | Left | HVLT delayed recall | -0.08 | 0.631 | 0.792 | 40 | 0.03 | 0.829 | 0.883 | 45 | -0.49 | 0.621 | 0.621 |
| PRC → pMTG_left | Reconstruction-retrieval | Left | HVLT delayed recall | -0.23 | 0.151 | 0.792 | 39 | 0.24 | 0.115 | 0.259 | 45 | -2.12 | 0.034 | 0.286 |
| HIP → PRC_right | Encoding-binding | Right | HVLT delayed recall | 0.08 | 0.608 | 0.780 | 40 | 0.07 | 0.661 | 0.680 | 45 | 0.07 | 0.941 | 0.941 |
| PRC → HIP_right | Encoding-binding | Right | HVLT delayed recall | 0.17 | 0.292 | 0.657 | 39 | -0.23 | 0.124 | 0.680 | 45 | 1.81 | 0.070 | 0.550 |
| VLPFC → HIP_right | Encoding-binding | Right | HVLT delayed recall | 0.17 | 0.301 | 0.657 | 39 | -0.14 | 0.392 | 0.680 | 41 | 1.33 | 0.183 | 0.550 |
| VLPFC → PRC_right | Encoding-binding | Right | HVLT delayed recall | -0.15 | 0.365 | 0.657 | 40 | -0.07 | 0.654 | 0.680 | 43 | -0.34 | 0.733 | 0.941 |
| AG → VLPFC_right | Reconstruction-retrieval | Right | HVLT delayed recall | -0.05 | 0.780 | 0.780 | 39 | -0.07 | 0.680 | 0.680 | 41 | 0.09 | 0.931 | 0.941 |
| HIP → AG_right | Reconstruction-retrieval | Right | HVLT delayed recall | -0.25 | 0.116 | 0.657 | 40 | -0.15 | 0.340 | 0.680 | 44 | -0.48 | 0.628 | 0.941 |
| HIP → pMTG_right | Reconstruction-retrieval | Right | HVLT delayed recall | 0.20 | 0.217 | 0.657 | 39 | -0.11 | 0.484 | 0.680 | 45 | 1.38 | 0.169 | 0.550 |
| pMTG → AG_right | Reconstruction-retrieval | Right | HVLT delayed recall | -0.05 | 0.756 | 0.780 | 39 | -0.13 | 0.410 | 0.680 | 45 | 0.33 | 0.740 | 0.941 |
| PRC → pMTG_right | Reconstruction-retrieval | Right | HVLT delayed recall | -0.10 | 0.522 | 0.780 | 40 | 0.09 | 0.531 | 0.680 | 46 | -0.89 | 0.373 | 0.840 |

*Table S10 continued …*

*Table S10 continued …*

| Directed connectivity | | | | Younger adults | | | | Older adults | | | | Younger vs older adults | | |
|---|---|---|---|---|---|---|---|---|---|---|---|---|---|---|
| Connection | Division | Hem | Measure | r | p | p-FDR | N | r | p | p-FDR | N | Z | p | p-FDR |
| Affective regulatory | | | | | | | | | | | | | | |
| ACC → AMY_left | Frontolimbic | Left | Beck anxiety | -0.08 | 0.620 | 0.775 | 40 | -0.05 | 0.740 | 0.926 | 44 | -0.13 | 0.896 | 0.994 |
| ACC → CAU_left | Frontolimbic | Left | Beck anxiety | 0.17 | 0.292 | 0.526 | 40 | -0.07 | 0.634 | 0.926 | 45 | 1.09 | 0.276 | 0.552 |
| AMY → DLPFC_left | Frontolimbic | Left | Beck anxiety | 0.26 | 0.108 | 0.360 | 39 | 0.00 | 0.987 | 0.987 | 44 | 1.16 | 0.246 | 0.552 |
| DLPFC → AMY_left | Frontolimbic | Left | Beck anxiety | 0.43 | 0.006 | 0.058 | 40 | 0.06 | 0.711 | 0.926 | 43 | 1.75 | 0.080 | 0.552 |
| DLPFC → CAU_left | Frontolimbic | Left | Beck anxiety | -0.05 | 0.741 | 0.775 | 39 | -0.05 | 0.730 | 0.926 | 45 | -0.01 | 0.994 | 0.994 |
| VLPFC → AMY_left | Frontolimbic | Left | Beck anxiety | 0.29 | 0.072 | 0.359 | 39 | 0.09 | 0.556 | 0.926 | 44 | 0.91 | 0.361 | 0.602 |
| ACC → aINS_left | Salience | Left | Beck anxiety | -0.21 | 0.202 | 0.506 | 39 | -0.16 | 0.303 | 0.926 | 45 | -0.24 | 0.813 | 0.994 |
| aINS → ACC_left | Salience | Left | Beck anxiety | -0.05 | 0.775 | 0.775 | 40 | -0.02 | 0.916 | 0.987 | 44 | -0.13 | 0.893 | 0.994 |
| aINS → VLPFC_left | Salience | Left | Beck anxiety | -0.06 | 0.722 | 0.775 | 39 | 0.27 | 0.070 | 0.695 | 45 | -1.49 | 0.135 | 0.552 |
| AMY → aINS_left | Salience | Left | Beck anxiety | 0.16 | 0.316 | 0.526 | 39 | -0.09 | 0.565 | 0.926 | 44 | 1.12 | 0.263 | 0.552 |
| ACC → AMY_right | Frontolimbic | Right | Beck anxiety | -0.02 | 0.908 | 0.908 | 39 | -0.10 | 0.538 | 0.673 | 44 | 0.34 | 0.737 | 0.819 |
| ACC → CAU_right | Frontolimbic | Right | Beck anxiety | 0.34 | 0.036 | 0.181 | 39 | 0.03 | 0.854 | 0.854 | 44 | 1.41 | 0.159 | 0.530 |
| AMY → DLPFC_right | Frontolimbic | Right | Beck anxiety | -0.20 | 0.210 | 0.390 | 40 | -0.06 | 0.688 | 0.765 | 45 | -0.64 | 0.523 | 0.771 |
| DLPFC → AMY_right | Frontolimbic | Right | Beck anxiety | 0.23 | 0.155 | 0.387 | 39 | 0.10 | 0.534 | 0.673 | 44 | 0.61 | 0.540 | 0.771 |
| DLPFC → CAU_right | Frontolimbic | Right | Beck anxiety | -0.20 | 0.234 | 0.390 | 39 | -0.18 | 0.253 | 0.673 | 44 | -0.09 | 0.931 | 0.931 |
| VLPFC → AMY_right | Frontolimbic | Right | Beck anxiety | 0.05 | 0.761 | 0.845 | 39 | 0.13 | 0.416 | 0.673 | 41 | -0.35 | 0.728 | 0.819 |
| ACC → aINS_right | Salience | Right | Beck anxiety | -0.06 | 0.713 | 0.845 | 40 | -0.21 | 0.161 | 0.673 | 45 | 0.69 | 0.489 | 0.771 |
| aINS → ACC_right | Salience | Right | Beck anxiety | 0.10 | 0.553 | 0.790 | 39 | -0.12 | 0.427 | 0.673 | 44 | 0.97 | 0.331 | 0.771 |
| aINS → VLPFC_right | Salience | Right | Beck anxiety | 0.43 | 0.006 | 0.059 | 39 | 0.11 | 0.479 | 0.673 | 42 | 1.52 | 0.129 | 0.530 |
| AMY → aINS_right | Salience | Right | Beck anxiety | 0.30 | 0.064 | 0.215 | 40 | -0.13 | 0.397 | 0.673 | 44 | 1.92 | 0.055 | 0.530 |
| ACC → AMY_left | Frontolimbic | Left | Beck depression | -0.20 | 0.224 | 0.447 | 40 | -0.07 | 0.670 | 0.917 | 44 | -0.59 | 0.557 | 0.717 |
| ACC → CAU_left | Frontolimbic | Left | Beck depression | 0.62 | 0.000 | 0.000 | 40 | -0.03 | 0.847 | 0.917 | 45 | 3.36 | 0.001 | 0.008 |
| AMY → DLPFC_left | Frontolimbic | Left | Beck depression | 0.09 | 0.573 | 0.650 | 39 | -0.09 | 0.574 | 0.917 | 44 | 0.79 | 0.429 | 0.717 |
| DLPFC → AMY_left | Frontolimbic | Left | Beck depression | 0.48 | 0.002 | 0.010 | 40 | 0.38 | 0.012 | 0.121 | 43 | 0.51 | 0.607 | 0.717 |
| DLPFC → CAU_left | Frontolimbic | Left | Beck depression | -0.08 | 0.610 | 0.650 | 39 | 0.02 | 0.897 | 0.917 | 45 | -0.46 | 0.646 | 0.717 |
| VLPFC → AMY_left | Frontolimbic | Left | Beck depression | 0.08 | 0.650 | 0.650 | 39 | 0.09 | 0.547 | 0.917 | 44 | -0.08 | 0.936 | 0.936 |
| ACC → aINS_left | Salience | Left | Beck depression | -0.31 | 0.057 | 0.191 | 39 | -0.16 | 0.299 | 0.917 | 45 | -0.69 | 0.488 | 0.717 |
| aINS → ACC_left | Salience | Left | Beck depression | 0.14 | 0.378 | 0.630 | 40 | -0.02 | 0.917 | 0.917 | 44 | 0.71 | 0.479 | 0.717 |
| aINS → VLPFC_left | Salience | Left | Beck depression | -0.08 | 0.623 | 0.650 | 39 | 0.11 | 0.475 | 0.917 | 45 | -0.84 | 0.400 | 0.717 |
| AMY → aINS_left | Salience | Left | Beck depression | 0.24 | 0.142 | 0.355 | 39 | -0.22 | 0.146 | 0.729 | 44 | 2.06 | 0.039 | 0.196 |
| ACC → AMY_right | Frontolimbic | Right | Beck depression | -0.09 | 0.568 | 0.876 | 39 | -0.11 | 0.464 | 0.931 | 44 | 0.08 | 0.933 | 0.933 |
| ACC → CAU_right | Frontolimbic | Right | Beck depression | 0.08 | 0.613 | 0.876 | 39 | -0.10 | 0.507 | 0.931 | 44 | 0.82 | 0.413 | 0.933 |
| AMY → DLPFC_right | Frontolimbic | Right | Beck depression | 0.02 | 0.881 | 0.978 | 40 | -0.08 | 0.609 | 0.931 | 45 | 0.46 | 0.648 | 0.933 |
| DLPFC → AMY_right | Frontolimbic | Right | Beck depression | 0.10 | 0.533 | 0.876 | 39 | 0.01 | 0.937 | 0.937 | 44 | 0.40 | 0.690 | 0.933 |
| DLPFC → CAU_right | Frontolimbic | Right | Beck depression | -0.15 | 0.347 | 0.876 | 39 | -0.09 | 0.578 | 0.931 | 44 | -0.30 | 0.760 | 0.933 |
| VLPFC → AMY_right | Frontolimbic | Right | Beck depression | 0.00 | 0.978 | 0.978 | 39 | -0.03 | 0.838 | 0.931 | 41 | 0.12 | 0.903 | 0.933 |
| ACC → aINS_right | Salience | Right | Beck depression | 0.11 | 0.517 | 0.876 | 40 | -0.22 | 0.139 | 0.931 | 45 | 1.48 | 0.139 | 0.933 |
| aINS → ACC_right | Salience | Right | Beck depression | 0.12 | 0.459 | 0.876 | 39 | 0.04 | 0.773 | 0.931 | 44 | 0.34 | 0.733 | 0.933 |
| aINS → VLPFC_right | Salience | Right | Beck depression | 0.04 | 0.820 | 0.978 | 39 | 0.13 | 0.394 | 0.931 | 42 | -0.42 | 0.671 | 0.933 |
| AMY → aINS_right | Salience | Right | Beck depression | 0.16 | 0.317 | 0.876 | 40 | 0.05 | 0.732 | 0.931 | 44 | 0.49 | 0.626 | 0.933 |

#### *4.5 Sensitivity analysis of Granger Causality lag parameter*

To evaluate the sensitivity of our vector autoregressive (VAR) frameworks and ensure that our primary brain-behaviour correlations were robust to model configurations, we conducted a systematic sensitivity analysis by recalculating the directed connectivity in each circuit across a range of model depths (Lags of 1, 3, and 5 fPET frames). Given our 16-second fPET frame resolution, these parameters represent distinct temporal windows spanning 16 seconds (Lag 1), 48 seconds (Lag 3), and 1 minute and 20 seconds (Lag 5), allowing us to assess how varying temporal boundaries alter the captured dynamics. At each alternative lag, individual subject Granger causality (GC) values were extracted and re-tested against the permutation null model to verify population-level network consistency, and subsequently correlated against all primary cognitive and affective measures. By explicitly assessing changes in model performance, parameter inflation, and behavioural sensitivity across these configurations, this sensitivity analysis serves to validate the empirical selection of Lag 2 (32s) as our primary analytical framework.

Across all evaluated functional domains, the multi-lag sensitivity analysis revealed a clear, parameter-dependent progression in both magnitude and statistical stability (Table S11). At the shortest interval (Lag 1; 16s), GC strengths heavily shrank toward zero and were largely unstable under permutation testing. Extending the model architecture to Lag 2 (32s) resolved a distinct, stable topography of directed flows, establishing prominent population-level significance across multiple core pathways (also see main manuscript). Lag 3 results were relatively consistent with Lag 2 in terms of significant pathways. Increasing the model depth Lag 5 (1 min 20s) introduced a systematic inflation of raw connectivity metrics, nearly tripling the baseline magnitudes. However, this also came at a cost: while Lag 5 captured select late, longer-window network connections (e.g., HIP to AG), it caused several primary population pathways (e.g., left ACC to CAU) to lose their statistical significance.

When evaluating brain-behaviour relationships, the choice of lag demonstrated an impact on behavioural sensitivity, revealing a shifting profile of cognitive and affective correlations (Table S12). At Lag 1 (16s), the framework captured a high density of correlations - particularly across bilateral frontolimbic and salience loops - but largely missed key memory and executive components. While Lag 3 maintained a profile of significant connections relatively comparable to Lag 2, moving to Lag 5 resulted in a notable reduction in the overall number of significant connectivity-behaviour associations, indicating a drop in sensitivity to individual differences..

Taken together, our multi-lag comparison highlights a trade-off between model underfitting and parameter-driven variance inflation in fPET effective connectivity. A 16-second window (Lag 1) underfits the slow metabolic dynamics of fPET, failing to capture stable population-level network topographies. Meanwhile, windows extending past one minute (Lags 5) overfit the data via parameter inflation, washing out brain-behaviour relationships and introducing statistical attrition across core cognitive and affective measures. Ultimately, Lag 2 (32 seconds) emerges as the optimal analytical window, minimising parameter inflation while preserving both population-level network features and maximum sensitivity to individual differences in cognitive performance and affective symptomatology.

**Table S11.** Sensitivity analysis of Granger Causality results to different lag parameters. Per the analyses in the main manuscript, the GC results for Lag 2 (32 seconds) are in the left columns and shaded blue. For comparison, sensitivity analysis was undertaken at Lag 1 (16s), 3 (48s) and 5 (1 min, 20 seconds).

| Circuit and connections | | | Lag 2 frames (32 seconds) | | | Lag 1 frame (16 seconds) | | | Lag 3 fames (48 seconds) | | | Lag 5 frames (1 min, 20 sec) | | |
|---|---|---|---|---|---|---|---|---|---|---|---|---|---|---|
| Connection | Hem | Circuit | Mean GC | SD | Null p | Mean GC | SD | Null p | Mean GC | SD | Null p | Mean GC | SD | Null p |
| DLPFC → ACC | Left | Cognitive control | 0.014 | 0.011 | 0.684 | 0.007 | 0.008 | 0.517 | 0.020 | 0.014 | 0.771 | 0.036 | 0.020 | 0.584 |
| DLPFC → CAU | Left | Cognitive control | 0.019 | 0.018 | 0.013 | 0.007 | 0.009 | 0.302 | 0.024 | 0.016 | 0.092 | 0.041 | 0.022 | 0.052 |
| ACC → CAU | Left | Cognitive control | 0.018 | 0.015 | 0.022 | 0.006 | 0.006 | 0.876 | 0.025 | 0.018 | 0.031 | 0.039 | 0.024 | 0.134 |
| ACC → DLPFC | Left | Cognitive control | 0.011 | 0.012 | 0.990 | 0.007 | 0.008 | 0.418 | 0.021 | 0.016 | 0.592 | 0.040 | 0.025 | 0.067 |
| VLPFC → PPC | Left | Cognitive control | 0.014 | 0.012 | 0.495 | 0.006 | 0.008 | 0.705 | 0.020 | 0.015 | 0.743 | 0.036 | 0.018 | 0.712 |
| aINS → ACC | Left | Cognitive control | 0.015 | 0.013 | 0.212 | 0.009 | 0.011 | 0.046 | 0.024 | 0.018 | 0.052 | 0.039 | 0.022 | 0.118 |
| aINS → VLPFC | Left | Cognitive control | 0.017 | 0.013 | 0.392 | 0.010 | 0.011 | 0.219 | 0.027 | 0.017 | 0.272 | 0.046 | 0.025 | 0.261 |
| VLPFC → IFJ | Left | Cognitive control | 0.014 | 0.010 | 0.568 | 0.007 | 0.008 | 0.371 | 0.023 | 0.016 | 0.208 | 0.041 | 0.024 | 0.064 |
| IFJ → PPC | Left | Cognitive control | 0.013 | 0.011 | 0.708 | 0.008 | 0.009 | 0.151 | 0.020 | 0.013 | 0.763 | 0.037 | 0.021 | 0.556 |
| DLPFC → ACC | Right | Cognitive control | 0.013 | 0.011 | 0.684 | 0.007 | 0.008 | 0.542 | 0.023 | 0.018 | 0.345 | 0.043 | 0.029 | 0.028 |
| DLPFC → CAU | Right | Cognitive control | 0.016 | 0.013 | 0.103 | 0.008 | 0.011 | 0.194 | 0.027 | 0.020 | 0.009 | 0.046 | 0.026 | 0.001 |
| ACC → CAU | Right | Cognitive control | 0.013 | 0.013 | 0.644 | 0.008 | 0.010 | 0.136 | 0.021 | 0.016 | 0.461 | 0.039 | 0.025 | 0.162 |
| ACC → DLPFC | Right | Cognitive control | 0.014 | 0.012 | 0.457 | 0.005 | 0.007 | 0.977 | 0.019 | 0.016 | 0.916 | 0.037 | 0.025 | 0.581 |
| VLPFC → PPC | Right | Cognitive control | 0.015 | 0.014 | 0.265 | 0.006 | 0.007 | 0.832 | 0.024 | 0.019 | 0.170 | 0.042 | 0.028 | 0.051 |
| aINS → ACC | Right | Cognitive control | 0.014 | 0.012 | 0.525 | 0.006 | 0.007 | 0.816 | 0.022 | 0.016 | 0.263 | 0.037 | 0.022 | 0.424 |
| aINS → VLPFC | Right | Cognitive control | 0.012 | 0.013 | 0.959 | 0.005 | 0.008 | 0.989 | 0.021 | 0.017 | 0.782 | 0.038 | 0.023 | 0.701 |
| IFJ → PPC | Right | Cognitive control | 0.019 | 0.016 | 0.004 | 0.007 | 0.009 | 0.365 | 0.024 | 0.017 | 0.119 | 0.043 | 0.027 | 0.048 |
| VLPFC → IFJ | Right | Cognitive control | 0.013 | 0.014 | 0.617 | 0.005 | 0.007 | 0.967 | 0.023 | 0.018 | 0.152 | 0.037 | 0.024 | 0.389 |
| IPS → DLPFC | Left | Working memory | 0.012 | 0.010 | 0.901 | 0.007 | 0.008 | 0.502 | 0.023 | 0.015 | 0.189 | 0.039 | 0.020 | 0.124 |
| DLPFC → IPS | Left | Working memory | 0.016 | 0.013 | 0.099 | 0.007 | 0.008 | 0.325 | 0.024 | 0.016 | 0.085 | 0.040 | 0.020 | 0.102 |
| DLPFC → VLPFC | Left | Working memory | 0.014 | 0.012 | 0.632 | 0.007 | 0.008 | 0.593 | 0.021 | 0.016 | 0.459 | 0.036 | 0.018 | 0.643 |
| mPFC → DLPFC | Left | Working memory | 0.014 | 0.013 | 0.364 | 0.008 | 0.008 | 0.241 | 0.021 | 0.013 | 0.566 | 0.040 | 0.024 | 0.073 |
| mPFC → IPL | Left | Working memory | 0.018 | 0.016 | 0.013 | 0.008 | 0.009 | 0.265 | 0.023 | 0.016 | 0.346 | 0.039 | 0.022 | 0.195 |
| VLPFC → IPL | Left | Working memory | 0.014 | 0.012 | 0.486 | 0.006 | 0.006 | 0.858 | 0.023 | 0.018 | 0.247 | 0.041 | 0.022 | 0.049 |
| IPL → VLPFC | Left | Working memory | 0.015 | 0.016 | 0.165 | 0.005 | 0.006 | 0.982 | 0.024 | 0.018 | 0.125 | 0.036 | 0.020 | 0.594 |
| IPL → IPS | Left | Working memory | 0.012 | 0.011 | 0.966 | 0.008 | 0.009 | 0.659 | 0.027 | 0.020 | 0.596 | 0.034 | 0.019 | 0.985 |
| IPS → DLPFC | Right | Working memory | 0.016 | 0.014 | 0.095 | 0.008 | 0.010 | 0.128 | 0.020 | 0.015 | 0.640 | 0.035 | 0.020 | 0.731 |
| DLPFC → IPS | Right | Working memory | 0.016 | 0.013 | 0.072 | 0.009 | 0.011 | 0.057 | 0.024 | 0.020 | 0.054 | 0.043 | 0.028 | 0.006 |
| DLPFC → VLPFC | Right | Working memory | 0.018 | 0.017 | 0.017 | 0.006 | 0.009 | 0.643 | 0.025 | 0.019 | 0.060 | 0.045 | 0.025 | 0.003 |
| mPFC → DLPFC | Right | Working memory | 0.013 | 0.012 | 0.629 | 0.006 | 0.006 | 0.885 | 0.021 | 0.014 | 0.618 | 0.037 | 0.023 | 0.479 |
| mPFC → IPL | Right | Working memory | 0.015 | 0.013 | 0.340 | 0.007 | 0.009 | 0.425 | 0.027 | 0.020 | 0.004 | 0.040 | 0.025 | 0.103 |
| VLPFC → IPL | Right | Working memory | 0.014 | 0.014 | 0.673 | 0.006 | 0.007 | 0.893 | 0.021 | 0.017 | 0.702 | 0.041 | 0.025 | 0.130 |
| IPL → VLPFC | Right | Working memory | 0.013 | 0.012 | 0.720 | 0.007 | 0.010 | 0.335 | 0.025 | 0.021 | 0.025 | 0.041 | 0.023 | 0.116 |
| IPL → IPS | Right | Working memory | 0.016 | 0.013 | 0.744 | 0.005 | 0.006 | 0.979 | 0.019 | 0.138 | 0.982 | 0.044 | 0.026 | 0.737 |
| VLPFC → PRC | Left | Episodic memory | 0.014 | 0.011 | 0.496 | 0.006 | 0.007 | 0.798 | 0.022 | 0.015 | 0.501 | 0.041 | 0.022 | 0.097 |
| VLPFC → HIP | Left | Episodic memory | 0.013 | 0.013 | 0.765 | 0.006 | 0.007 | 0.862 | 0.021 | 0.016 | 0.464 | 0.035 | 0.019 | 0.722 |
| PRC → HIP | Left | Episodic memory | 0.014 | 0.014 | 0.502 | 0.005 | 0.006 | 0.955 | 0.023 | 0.017 | 0.259 | 0.035 | 0.023 | 0.627 |
| HIP → PRC | Left | Episodic memory | 0.014 | 0.013 | 0.526 | 0.005 | 0.005 | 0.989 | 0.022 | 0.016 | 0.433 | 0.038 | 0.024 | 0.251 |
| PRC → pMTG | Left | Episodic memory | 0.014 | 0.013 | 0.476 | 0.006 | 0.008 | 0.781 | 0.024 | 0.017 | 0.110 | 0.044 | 0.027 | 0.011 |
| HIP → pMTG | Left | Episodic memory | 0.015 | 0.014 | 0.272 | 0.007 | 0.008 | 0.602 | 0.025 | 0.018 | 0.036 | 0.039 | 0.021 | 0.187 |
| pMTG → AG | Left | Episodic memory | 0.014 | 0.012 | 0.455 | 0.006 | 0.007 | 0.869 | 0.021 | 0.016 | 0.543 | 0.037 | 0.021 | 0.445 |
| HIP → AG | Left | Episodic memory | 0.015 | 0.014 | 0.229 | 0.007 | 0.008 | 0.600 | 0.024 | 0.018 | 0.111 | 0.045 | 0.028 | 0.003 |
| AG → VLPFC | Left | Episodic memory | 0.016 | 0.014 | 0.077 | 0.008 | 0.011 | 0.058 | 0.023 | 0.018 | 0.133 | 0.036 | 0.019 | 0.571 |
| VLPFC → PRC | Right | Episodic memory | 0.017 | 0.015 | 0.022 | 0.008 | 0.009 | 0.122 | 0.026 | 0.018 | 0.018 | 0.042 | 0.024 | 0.028 |
| VLPFC → HIP | Right | Episodic memory | 0.012 | 0.011 | 0.909 | 0.006 | 0.009 | 0.724 | 0.019 | 0.016 | 0.862 | 0.037 | 0.023 | 0.329 |
| PRC → HIP | Right | Episodic memory | 0.012 | 0.012 | 0.937 | 0.006 | 0.008 | 0.709 | 0.020 | 0.017 | 0.816 | 0.037 | 0.026 | 0.532 |
| HIP → PRC | Right | Episodic memory | 0.014 | 0.012 | 0.492 | 0.007 | 0.009 | 0.300 | 0.020 | 0.015 | 0.678 | 0.041 | 0.026 | 0.159 |
| PRC → pMTG | Right | Episodic memory | 0.016 | 0.014 | 0.119 | 0.006 | 0.008 | 0.631 | 0.024 | 0.018 | 0.080 | 0.040 | 0.025 | 0.109 |
| HIP → pMTG | Right | Episodic memory | 0.014 | 0.012 | 0.443 | 0.007 | 0.009 | 0.485 | 0.023 | 0.016 | 0.171 | 0.036 | 0.023 | 0.586 |
| pMTG → AG | Right | Episodic memory | 0.012 | 0.010 | 0.886 | 0.009 | 0.011 | 0.044 | 0.023 | 0.019 | 0.292 | 0.036 | 0.022 | 0.570 |
| HIP → AG | Right | Episodic memory | 0.013 | 0.011 | 0.691 | 0.006 | 0.007 | 0.861 | 0.022 | 0.018 | 0.298 | 0.039 | 0.025 | 0.233 |
| AG → VLPFC | Right | Episodic memory | 0.015 | 0.015 | 0.348 | 0.006 | 0.008 | 0.726 | 0.021 | 0.017 | 0.528 | 0.037 | 0.022 | 0.394 |
| DLPFC → AMY | Left | Affective regulatory | 0.014 | 0.012 | 0.421 | 0.008 | 0.011 | 0.112 | 0.023 | 0.016 | 0.179 | 0.039 | 0.024 | 0.216 |
| VLPFC → AMY | Left | Affective regulatory | 0.012 | 0.011 | 0.889 | 0.006 | 0.007 | 0.725 | 0.024 | 0.016 | 0.083 | 0.039 | 0.022 | 0.189 |
| AMY → DLPFC | Left | Affective regulatory | 0.015 | 0.013 | 0.180 | 0.008 | 0.008 | 0.103 | 0.024 | 0.016 | 0.134 | 0.037 | 0.021 | 0.410 |
| ACC → CAU | Left | Affective regulatory | 0.018 | 0.015 | 0.022 | 0.006 | 0.006 | 0.876 | 0.025 | 0.018 | 0.031 | 0.039 | 0.024 | 0.134 |
| ACC → AMY | Left | Affective regulatory | 0.013 | 0.011 | 0.759 | 0.007 | 0.007 | 0.631 | 0.021 | 0.014 | 0.597 | 0.039 | 0.018 | 0.189 |
| DLPFC → CAU | Left | Affective regulatory | 0.019 | 0.018 | 0.013 | 0.007 | 0.009 | 0.302 | 0.024 | 0.016 | 0.092 | 0.041 | 0.022 | 0.052 |
| AMY → aINS | Left | Affective regulatory | 0.013 | 0.010 | 0.721 | 0.005 | 0.005 | 0.908 | 0.021 | 0.016 | 0.481 | 0.042 | 0.024 | 0.010 |
| ACC → aINS | Left | Affective regulatory | 0.017 | 0.013 | 0.042 | 0.008 | 0.008 | 0.245 | 0.025 | 0.018 | 0.028 | 0.039 | 0.021 | 0.127 |
| aINS → VLPFC | Left | Affective regulatory | 0.017 | 0.013 | 0.392 | 0.010 | 0.011 | 0.219 | 0.027 | 0.017 | 0.272 | 0.046 | 0.025 | 0.261 |
| aINS → ACC | Left | Affective regulatory | 0.015 | 0.013 | 0.212 | 0.009 | 0.011 | 0.046 | 0.024 | 0.018 | 0.052 | 0.039 | 0.022 | 0.118 |
| AMY → DLPFC | Right | Affective regulatory | 0.015 | 0.012 | 0.192 | 0.007 | 0.008 | 0.447 | 0.022 | 0.017 | 0.299 | 0.037 | 0.022 | 0.399 |
| DLPFC → CAU | Right | Affective regulatory | 0.016 | 0.013 | 0.103 | 0.008 | 0.011 | 0.194 | 0.027 | 0.020 | 0.009 | 0.046 | 0.026 | 0.001 |
| VLPFC → AMY | Right | Affective regulatory | 0.014 | 0.012 | 0.466 | 0.006 | 0.008 | 0.602 | 0.019 | 0.015 | 0.835 | 0.036 | 0.022 | 0.533 |
| ACC → AMY | Right | Affective regulatory | 0.014 | 0.012 | 0.450 | 0.006 | 0.008 | 0.789 | 0.021 | 0.014 | 0.604 | 0.037 | 0.020 | 0.435 |
| ACC → CAU | Right | Affective regulatory | 0.013 | 0.013 | 0.644 | 0.008 | 0.010 | 0.136 | 0.021 | 0.016 | 0.461 | 0.039 | 0.025 | 0.162 |
| DLPFC → AMY | Right | Affective regulatory | 0.013 | 0.014 | 0.730 | 0.005 | 0.009 | 0.932 | 0.021 | 0.019 | 0.498 | 0.039 | 0.027 | 0.154 |
| aINS → ACC | Right | Affective regulatory | 0.014 | 0.012 | 0.525 | 0.006 | 0.007 | 0.816 | 0.022 | 0.016 | 0.263 | 0.037 | 0.022 | 0.424 |
| AMY → aINS | Right | Affective regulatory | 0.016 | 0.015 | 0.117 | 0.007 | 0.010 | 0.390 | 0.023 | 0.017 | 0.165 | 0.041 | 0.023 | 0.061 |
| ACC → aINS | Right | Affective regulatory | 0.014 | 0.014 | 0.496 | 0.007 | 0.010 | 0.400 | 0.022 | 0.016 | 0.347 | 0.041 | 0.024 | 0.044 |
| aINS → VLPFC | Right | Affective regulatory | 0.012 | 0.013 | 0.959 | 0.005 | 0.008 | 0.989 | 0.021 | 0.017 | 0.782 | 0.038 | 0.023 | 0.701 |

**Table S12.** Sensitivity analysis of Granger Causality (GC) calculated at different Lags (number of fPET frames), showing significant correlations with cognition and psychosocial function measures at $p < 0.05$. Per the analyses in the main manuscript, the GC results for Lag 2 (32 seconds) are shown at the top of the table shaded blue. Sensitivity analysis was undertaken at Lag 1 (16s), 3 (48s) and 5 (1 min, 20 seconds).

| Connection | Division | Hem | Measure | Correlation | P-Value |
|---|---|---|---|---|---|
| Lag 2 frames (32seconds; see main maunscript) | | | | | |
| DLPFC → ACC | Proactive | Right | Switch cost (reversed) | 0.325 | 0.002 |
| DLPFC → CAU | Proactive | Right | Switch cost (reversed) | 0.221 | 0.044 |
| aINS → ACC | Reactive | Right | Stop signal RT (reversed) | 0.304 | 0.005 |
| VLPFC → IFJ | Reactive | Right | Stop signal RT (reversed) | -0.309 | 0.005 |
| HIP → PRC | Encoding-binding | Left | HVLT delayed recall | 0.291 | 0.007 |
| IPL → IPS | Storage-maintenance | Left | Digit span total | 0.217 | 0.047 |
| DLPFC → IPS | Attention-control | Right | Digit span total | -0.290 | 0.007 |
| DLPFC → AMY | Frontolimbic | Left | Beck depression | 0.461 | 0.000 |
| DLPFC → AMY | Frontolimbic | Left | Beck anxiety | 0.338 | 0.002 |
| ACC → CAU | Frontolimbic | Left | Beck depression | 0.264 | 0.015 |
| ACC → aINS | Salience | Left | Beck depression | -0.242 | 0.027 |
| aINS → VLPFC | Salience | Right | Beck anxiety | 0.411 | 0.000 |
| Lag 1 frame (16 seconds) | | | | | |
| Connection | Division | Hem | Measure | Correlation | P-Value |
| DLPFC → ACC | Proactive | Right | Switch cost (reversed) | 0.291 | 0.007 |
| ACC → DLPFC | Proactive | Right | Switch cost (reversed) | -0.278 | 0.011 |
| IFJ → PPC | Reactive | Right | Stop signal RT (reversed) | 0.291 | 0.007 |
| DLPFC → VLPFC | Attention-control | Right | Digit span total | -0.260 | 0.020 |
| DLPFC → AMY | Frontolimbic | Left | Beck depression | 0.484 | 0.000 |
| VLPFC → AMY | Frontolimbic | Left | Beck anxiety | 0.270 | 0.014 |
| ACC → AMY | Frontolimbic | Left | Beck depression | -0.228 | 0.037 |
| DLPFC → AMY | Frontolimbic | Right | Beck depression | 0.284 | 0.009 |
| DLPFC → AMY | Frontolimbic | Right | Beck anxiety | 0.413 | 0.000 |
| ACC → CAU | Frontolimbic | Right | Beck anxiety | 0.300 | 0.006 |
| DLPFC → CAU | Frontolimbic | Right | Beck depression | -0.223 | 0.043 |
| AMY → aINS | Salience | Right | Beck anxiety | 0.300 | 0.006 |
| Lag 3 frames (48 seconds) | | | | | |
| Connection | Division | Hem | Measure | Correlation | P-Value |
| ACC → CAU | Proactive / Frontolimbic | Left | Stop signal RT (reversed) | 0.251 | 0.020 |
| DLPFC → ACC | Proactive | Right | Switch cost (reversed) | 0.240 | 0.027 |
| VLPFC → PPC | Reactive | Right | Switch cost (reversed) | 0.225 | 0.044 |
| HIP → PRC | Encoding-binding | Left | HVLT delayed recall | 0.263 | 0.014 |
| DLPFC → IPS | Attention-control | Right | Digit span total | -0.332 | 0.002 |
| DLPFC → AMY | Frontolimbic | Left | Beck depression | 0.388 | 0.000 |
| ACC → CAU | Frontolimbic | Left | Beck depression | 0.388 | 0.000 |
| DLPFC → AMY | Frontolimbic | Right | Beck anxiety | 0.445 | 0.000 |
| aINS → ACC | Salience | Right | Beck depression | 0.284 | 0.009 |
| aINS → VLPFC | Salience | Right | Beck anxiety | 0.237 | 0.033 |
| Lag 5 frames (1 minute 20 seconds) | | | | | |
| Connection | Division | Hem | Measure | Correlation | P-Value |
| ACC → CAU | Proactive / Frontolimbic | Left | Stop signal RT (reversed) | 0.252 | 0.020 |
| VLPFC → PPC | Reactive | Right | Switch cost (reversed) | 0.272 | 0.013 |
| DLPFC → AMY | Frontolimbic | Left | Beck depression | 0.345 | 0.001 |
| DLPFC → AMY | Frontolimbic | Right | Beck anxiety | 0.366 | 0.001 |
| aINS → ACC | Salience | Right | Beck depression | 0.220 | 0.046 |
